# Relative contributions of correcting the diet and voluntary exercise to myocardial recovery in a two-hit murine model of heart failure with preserved ejection fraction

**DOI:** 10.1101/2025.02.03.636256

**Authors:** Emylie-Ann Labbé, Sara-Ève Thibodeau, Élisabeth Walsh-Wilkinson, Maude Chalifour, Pierre-Olivier Sirois, Juliette Leblanc, Marie Arsenault, Jacques Couet

**Affiliations:** Groupe de recherche sur les valvulopathies, Centre de recherche de l’Institut universitaire de cardiologie et de pneumologie de Québec, Université Laval, Québec City, Québec, Canada

**Keywords:** mouse, heart failure, myocardial recovery, reverse remodelling, cardiac hypertrophy, life habits, preclinical model, HFpEF

## Abstract

We recently proposed a new two-hit murine model of heart failure with preserved ejection fraction (HFpEF) amenable for both male and female animals. Using this model, we studied cardiac reverse remodelling (RR) after stopping the causing stress (Angiotensin II (AngII) + High-fat diet; MHS) after 28 days, and then introducing voluntary exercise (VE) and feeding the animals with a low-fat diet. We showed this could lead to extensive cardiac left ventricle (LV) RR. Revisiting this HFpEF model, we studied the relative contribution to RR of correcting the diet and/or starting VE after stopping AngII. We also evaluate the extent of myocardial recovery after an extended period (12 weeks instead of four) by exposing the animals to a second MHS.

Our observations revealed a sex-specific response. Discontinuing AngII but continuing the HFD blocked RR in females, not males. Removing AngII and correcting either the diet or implementing VE normalized most tested gene markers of LV hypertrophy or extracellular matrix, irrespective of sex.

Twelve weeks of recovery was associated with normal LV morphology and function, except for abnormal diastolic echocardiographic parameters. A second MHS after these 12 weeks led to a loss of ejection fraction in males and LV dilatation. The response of females was like that after the first MHS, suggesting a better recovery. The MHS changed markers of myocardial glucose metabolism. Pyruvate dehydrogenase (PDH) activity responsible for pyruvate entry in the mitochondria was reduced after MHS, and this was accompanied by an increase of PDH phosphorylation and pyruvate dehydrogenase kinase 4 content. RR mostly normalized these.

Our results suggest sex-specific RR after stopping the MHS and that myocardial anomalies remaining in males make them more sensitive to a second HFpEF-inducing stress.

## Introduction

Several clinical trials were conducted in patients with stable heart failure with preserved ejection fraction (HFpEF), showing that supervised exercise training was safe and could significantly improve exercise capacity and quality of life. Cardiac effects were mainly neutral, and benefits on clinical outcomes were limited (1). Changes in lifestyle combining physical activity and nutritional modifications also showed benefits in HFpEF patients (2). It is unclear whether these interventions, incorporating physical activity and caloric restriction, improve cardiac function or if the effects are mainly limited to peripheral targets, especially the skeletal muscle (3).

We recently observed in a “two-hit” murine HFpEF model (Angiotensin II (AngII) + High-Fat Diet (HFD) or MHS) that stopping AngII, normalizing the diet, and introducing voluntary exercise (VE) helped reverse cardiac damage and improved exercise capacity (4). It was unclear if correcting the diet or introducing VE contributed to the left ventricle (LV) reverse remodelling (RR) in addition to removing the hypertensive stressor, AngII.

Left ventricle RR has received limited attention in the context of HFpEF preclinical studies. Diet normalization in Dahl Salt-Sensitive helped reduce diastolic dysfunction, increased exercise capacity and lowered pulmonary congestion without effects on blood pressure and cardiac hypertrophy (5).

Another issue is the extent of myocardial recovery following LV reverse remodelling. Is normalizing various morphologic, functional, metabolic, and molecular parameters a good marker of myocardial recovery? Myocardial recovery in patients has been mainly studied in the context of heart failure with reduced ejection fraction or HFrEF (6). There are still uncertainties about the clinical conduct of patients who recovered ejection fraction and the appropriate markers of this recovery. Interestingly, data suggest that age, biological sex, etiology, duration of disease, and comorbidities are important variables associated with the likelihood of ejection fraction recovery (7).

LV reverse remodelling in patients with an HFpEF-like disease, such as aortic valve stenosis undergoing valve replacement, is usually present and is accompanied by hypertrophy regression. Myocardial fibrosis reduction is often present but happens later after valve replacement. Less than 20% of patients return to average LV mass after 18 months (8, 9).

To study the extent of myocardial recovery in our murine model, we investigated cardiac resiliency to a second MHS after an extended period of remission of 12 weeks instead of the four we already studied. We also investigated the relative contribution of physical activity and diet to RR.

A second MHS after a 12-week remission induces a more severe phenotype in males than in female mice. This phenotype is associated with eccentric LV remodelling and a loss of ejection fraction. These changes are related to corresponding modifications in the myocardial glucose metabolism. We also identified sex differences in HFpEF mice to the response to the diet during RR.

## Materials and Methods

### Animals

C57BL6/J male and female 7-week-old mice were purchased from Jackson Laboratory (Bar Harbor, ME, USA). Mice were housed on a 12h light-12h dark cycle with free access to chow and water. The protocol was approved by the Université Laval’s animal protection committee and followed the recommendations of the Canadian Council on Laboratory Animal Care (#2023-1249 and #2023-1250). This study was conducted in accordance with the ARRIVE guidelines. Mice were randomly distributed in the various experimental groups. Group size varied from 8 to 12 animals per group.

### Metabolic and hypertensive stress (MHS)

Males (M) or females (F) were implanted or not with an osmotic minipump (Alzet #1004) providing a continuous infusion of angiotensin II (AngII; 1.5mg/kg/day) (Sigma) for 28 days and fed (MHS or metabolic-hypertensive stress) or not with a high-fat diet (HFD: 60% calories; Research Diets Cat. #D12492). The low-fat diet (KFD) had 10% of calories from fat (D12450J) (4).

### Reverse remodelling

On day 28, osmotic minipumps were removed. Mice were then divided into three groups: 1) **RR**: the diet of the MHS mice was changed to the LFD, and the same day, a flying saucer-type exercise running device was introduced in the cages (Innowheel™, Innovive, Billerica, MA, USA) to allow voluntary exercise (VE), 2) **Sed**: the HFD was changed to the LFD diet, no VE, and 3) **HFD**: the HFD was continued, but VE was allowed. Twenty-eight days later, the protocol was stopped. A schematic representation of the study is illustrated in Figure 1A. Mice had an echocardiography (Echo) exam the day before sacrifice. Euthanasia and tissue collection were performed as previously described (4, 10)

**Figure 1.**
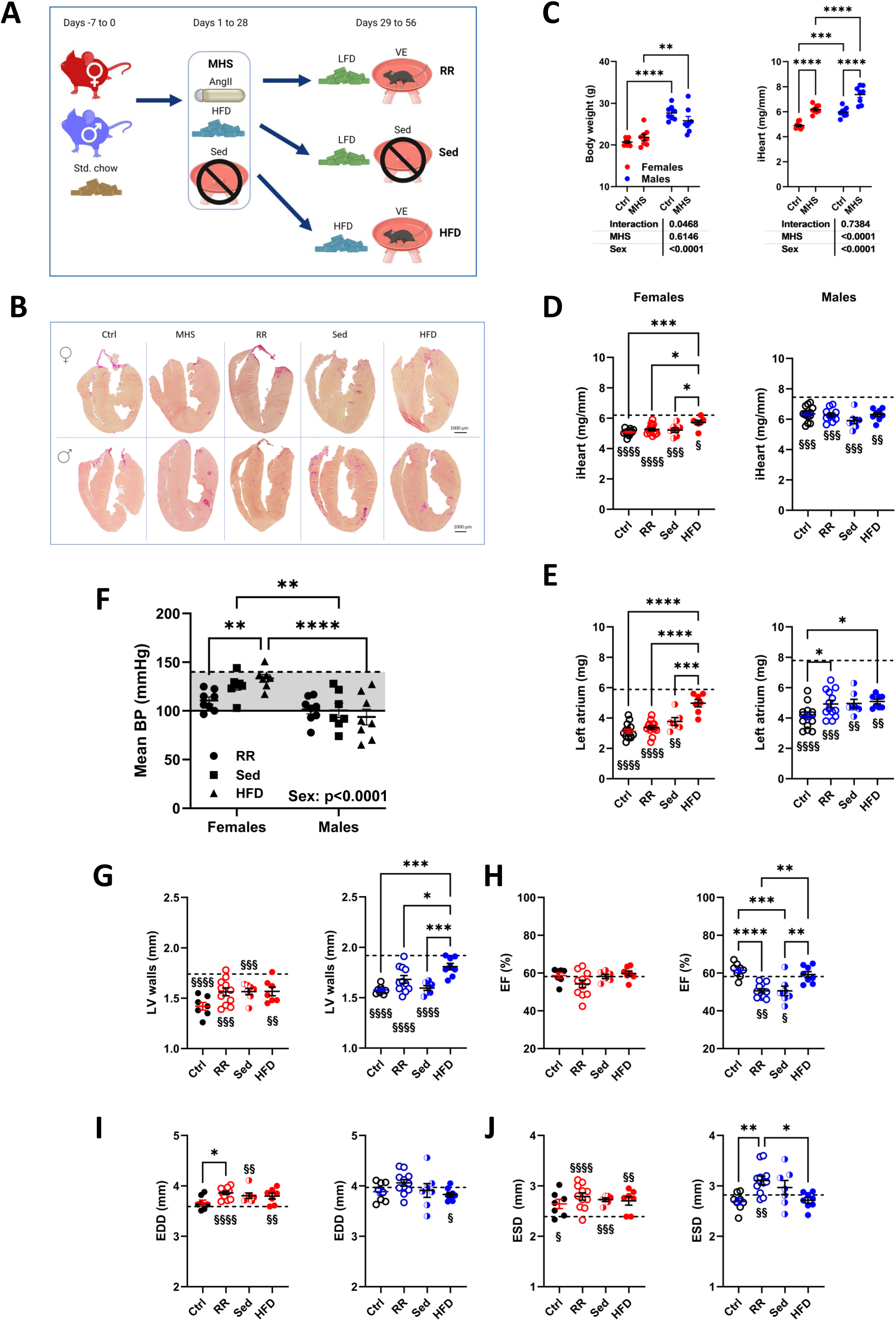
Continuing the high-fat diet (HFD) in female mice after the MHS alters cardiac reverse remodelling. A. Schematic representation of the experimental design (Created in BioRender. Couet, J. (2025) https://BioRender.com/j31s771). The weeks before the experiment, the mice were fed a standard diet (17% calories from fat). From day 1, males or females were implanted subcutaneously in their back, an osmotic minipump providing a continuous infusion of angiotensin II (AngII; 1.5mg/kg/day) for 28 days and fed with a high-fat diet (HFD) (**MHS**). On day 28, the AngII-releasing minipump was removed. Mice were then divided into three groups: 1) **RR**, the HFD was changed to the low-fat diet (LFD), and the same day, a flying saucer-type exercise device was introduced in their cage. 2) **Sed**, the HFD was changed to the LFD diet, but no exercise device was introduced in their cage; and 3) **HFD**, the HFD was continued, but voluntary exercise (VE) was allowed. Twenty-eight days later, the protocol was stopped. Control (Ctrl) mice were kept on the standard diet for the first 28 days and then were shifted to the LFD diet and VE. B. Representative images of picrosirius red staining of female and male heart sections for each indicated group. C. Body weight and indexed heart weight (iHeart) of female (red) and male (blue) mice after the MHS. Two-way ANOVA followed by Holm-Sidak post-test. *: p<0.05, **: p<0.01, ***: p<0.001 and ****: p<0.0001 between indicated groups. D. Indexed heart weight of mice after four weeks of recovery post-MHS. Ctrl: control, RR: reverse remodelling group, Sed: LFD without voluntary exercise (VE) and HFD: HFD and VE. E. Left atrial weight. The dotted line on a graph represents the mean of the indicated parameter after 28 days of MHS. F. Mean blood pressure in conscious mice after 28 days of recovery. The solid line represents the mean of the parameter in control mice and the dotted line after 28 days of MHS. G. LV wall thickness in diastole (PWd+IVSd). H. Ejection fraction (EF). I. End-diastolic LV diameter (EDD). J. End-systolic LV diameter (ESD). One-way ANOVA followed by Holm-Sidak post-test. *: p<0.05, **: p<0.01, ***: p<0.001 and ****: p<0.0001 between indicated groups. §: p<0.05, §§: p<0.01, §§§: p<0.001 and §§§§: p<0.0001 between the indicated group and MHS group using the Student’s T-test.

### Extended myocardial recovery (MR) and second MHS

Three additional sets of mice were studied. Male and female mice (8-week-old) were divided into Controls, MR, and MR + MHS. Controls: Mice were kept sedentary until 12 weeks of age on a standard diet (17% of calories from fat), then the animals were allowed VE for an additional 12 weeks and fed the LFD mimicking the RR regimen described above (Ctrl/VE12). MR: MHS for 28 days, then RR for 12 weeks. MR + MHS: After RR for 12 weeks, a second 28-day MHS was performed.

Experienced technicians monitored the mice’s health and behaviour daily during the protocol, and the animals were weighed weekly. Three male mice died in the MR + MHS group during the second MHS stress.

### Echocardiography

Echocardiography was performed under isoflurane anesthesia as described previously (10, 11).

### Measurements of Blood Pressure

As previously described, blood pressure was measured using the non-invasive CODA tail-cuff system (Kent Scientific, Torrington, CT) (4, 12).

### Myocardial Fibrosis Evaluation

Myocardial samples were fixated, sliced in serial sections (10μm thick), and stained with Picrosirius Red to assess the cross-sectional area and the percentage of interstitial fibrosis. The formula used to calculate the percentage of interstitial fibrosis was ((% Fibrosis) / (%Fibrosis + %Tissue)) * 100.

### Cardiomyocyte Cross-Sectional Area

The cardiomyocyte cross-sectional area (CSA) was visualized with immunofluorescent wheat germ agglutinin FITC (Sigma) staining. CSA was then measured and expressed in µm^2^ as previously described (12).

### RNA Isolation and Quantitative Real-Time Polymerase Chain Reaction

As described previously, total RNA from LV tissue was extracted (17). Quantitative RT-PCR quantified LV gene expression for at least six animals per group. Pre-optimized primers were from QuantiTect (Qiagen) and IDT (Coralville, Iowa), and Sso Advanced Universal SYBR Green Supermix (Bio-Rad, Hercules, CA) was used. Cyclophilin A (Ppia) was the control “housekeeping” gene. The primers used are listed in Table S1.

### Myocardial metabolism enzymatic activity

Citrate synthase (CS) and Hydroxyacyl-Coenzyme A dehydrogenase (HADH) myocardial enzymatic activities were measured as previously described on frozen pieces of mouse LV (∼10mg) (13). Following the supplier’s instructions, Pyruvate Dehydrogenase activity was assayed using an enzyme activity microplate assay kit from Abcam (#ab109902).

### Western blots

Protein content was estimated by Western blotting as described previously (14, 15). Antibodies were diluted 1:1000 in a TSB-T solution with 5% bovine serum albumin (BSA). Phospho-PDH was obtained from Cell Signaling Technologies (Danvers MA; #31866), whereas PDH (AB110416-1002) and PDK4 antibodies were from Abcam (AB214938) (Toronto, ON, Canada). The appropriate second antibody was diluted 1:2000 in TBS-T with 5% milk. Western Lightning Plus ECL (Perkin-Elmer, Woodbridge, ON, Canada) and Clarity Max ECL (Bio-Rad) were used for membrane revelation with the Gel Doc system and Image Lab software (Bio-Rad). The “Total lane method” was used for normalization, and adjustments were made according to pre-blocking membrane exposure and a standard pool run on each gel (16).

### Statistical Analysis

All data are expressed as mean ± standard error of the mean (SEM). Intergroup comparisons were conducted using the Student’s T-test using GraphPad Prism 10.4 (GraphPad Software Inc., La Jolla, CA, USA). Outliers were removed using the ROUT test with a Q of 1% with Prism.

Comparisons of more than two groups were analyzed using one-way or two-way ANOVA and Holm-Sidak post-test. P<0.05 was considered statistically significant.

## Results

### The Angiotensin II/High-Fat Diet stress (MHS) induces sex-specific changes in left ventricle global strain and diastolic function

Sixteen control mice and forty MHS mice (Figure 1A) of each sex had a complete echocardiographic exam at the end of the four weeks of the stress at the age of three months. The MHS mice showed thicker LV walls in diastole and systole (Table 1). End-diastolic and end-systolic LV diameters were reduced, but LV volumes remained unchanged. Ejection fraction and cardiac output also remained unchanged by the MHS. Global longitudinal and global circumference strains were significantly decreased in male MHS mice but not in females (Table 1). As described in Table 2, the left atrial diameter was increased by MHS. Both E and A waves were reduced after MHS in male animals but not females. Tissue Doppler E’ and A’ waves were both reduced in males but only the E’ wave in females. This resulted in an abnormal E/E’ ratio in females.

**Table 1.** Echo data in female and male mice after MHS. Standard echo left ventricle parameters after 4 weeks of MHS and after 4 weeks. Control mice were studied in parallel Echo exams as described in the Methods section. PWd: diastolic posterior wall thickness, IVSd: diastolic inter-ventricular septum, PWs: systolic posterior wall thickness, IVSs: systolic inter-ventricular septum, EDD: end-diastolic LV diameter, ESV: end-systolic LV diameter, RWT: relative wall thickness, EDV: end-diastolic volume, ESV: end-systolic volume, SV stroke volume, HR: heart rate, EF: ejection fraction, CO: cardiac output, GLS: global longitudinal strain and GCS: global circumferential strain. Results are expressed as the mean ± standard error of the mean (SEM). P values were calculated using the Student’s T-test.

| **Females** |  |  |  |
| --- | --- | --- | --- |
| Parameters | **Ctrl (n=16)** | **MHS (n=40)** | **P value** |
| PWd, mm | 0.71±0.013 | 0.91±0.011 | <0.0001 |
| IVSd, mm | 0.68±0.013 | 0.85±0.012 | <0.0001 |
| PWs, mm | 1.04±0.029 | 1.28±0.018 | <0.0001 |
| IVSs, mm | 0.94±0.017 | 1.10±0.017 | <0.0001 |
| EDD, mm | 3.69±0.050 | 3.49±0.033 | 0.0016 |
| ESD, mm | 2.54±0.059 | 2.32±0.041 | 0.0056 |
| RWT | 0.38±0.010 | 0.51±0.009 | <0.0001 |
| LV mass, mg | 84.6±1.67 | 108.2±2.16 | <0.0001 |
| EDV, μl | 46.7±1.64 | 43.6±1.45 | 0.23 |
| ESV, μl | 17.5±0.81 | 16.9±0.72 | 0.62 |
| SV, mm | 29.1±0.95 | 26.7±0.88 | 0.12 |
| HR, bpm | 510±10.7 | 515±6.5 | 0.72 |
| EF, % | 62.6±0.73 | 61.5±0.79 | 0.42 |
| CO, ml/min | 14.8±0.53 | 13.7±0.46 | 0.17 |
| GLS | -16.8±0.50 | -15.9±0.56 | 0.30 |
| GCS | -26.1±0.66 | -27.4±0.88 | 0.31 |
| **Males** |  |  |  |
| Parameters | **Ctrl (n=16)** | **MHS (n=40)** | **P value** |
| PWd, mm | 0.75±0.011 | 0.99±0.013 | <0.0001 |
| IVSd, mm | 0.75±0.008 | 0.89±0.009 | <0.0001 |
| PWs, mm | 1.06±0.020 | 1.30±0.016 | <0.0001 |
| IVSs, mm | 0.99±0.022 | 1.13±0.012 | <0.0001 |
| EDD, mm | 3.97±0.032 | 3.66±0.054 | 0.0007 |
| ESD, mm | 2.85±0.045 | 2.65±0.010 | 0.025 |
| RWT | 0.38±0.006 | 0.52±1.4 | <0.0001 |
| LV mass, mg | 106.7±1.84 | 128.0±3.05 | <0.0001 |
| EDV, μl | 52.8±2.10 | 50.4±2.04 | 0.50 |
| ESV, μl | 22.2±0.83 | 21.7±1.07 | 0.78 |
| SV, mm | 30.6±1.44 | 28.7±1.07 | 0.34 |
| HR, bpm | 489±10.5 | 525±5.4 | 0.0015 |
| EF, % | 57.8±0.88 | 57.3±0.81 | 0.71 |
| CO, ml/min | 14.9±0.71 | 15.1±0.59 | 0.90 |
| GLS | -16.7±0.53 | -14.4±0.73 | 0.031 |
| GCS | -27.7±0.85 | -24.3±1.04 | 0.048 |

**Table 2.** Echo diastolic parameters in female and male mice after MHS. Standard echo left ventricle parameters were measured in male and female mice. Echo exams were performed as described in the Methods section. LA diam: left atrial diameter, E: E wave. A: A wave, E slope: E wave slope, IVRT: isovolumetric relaxation time, IVCT: isovolumetric contraction time, MPI: myocardial performance index, E’: E’ wave, A’: A’ wave. Results are expressed as the mean ± standard error of the mean (SEM). P values were calculated using the Student’s T-test.

| **Females** |  |  |  |
| --- | --- | --- | --- |
| Parameters | **Ctrl (n=16)** | **MHS (n=40)** | **P value** |
| LA diam., mm | 2.09±0.024 | 2.32±0.026 | <0.0001 |
| E, mm/s | 571±15.7 | 575±10.5 | 0.85 |
| A, mm/s | 366±8.3 | 390±7.9 | 0.081 |
| E slope | -39859±1393.2 | -34510±1059.9 | 0.0074 |
| E/A | 1.57±0.034 | 1.48±0.024 | 0.070 |
| IVRT, ms | 15.5±0.23 | 16.1±0.20 | 0.096 |
| IVCT, ms | 12.1±0.22 | 11.5±0.14 | 0.063 |
| MPI | 0.68±0.015 | 0.68±0.011 | 0.88 |
| E', mm/s | -30.8±0.58 | -25.4±0.54 | <0.0001 |
| A', mm/s | -18.1±0.49 | -16.9±0.34 | 0.060 |
| E'/A' | 1.71±0.029 | 1.51±0.034 | 0.0015 |
| E/E' | -18.6±0.54 | -22.9±0.49 | <0.0001 |
| **Males** |  |  |  |
| Parameters | **Ctrl (n=16)** | **MHS (n=40)** | **P value** |
| LA diam., mm | 2.30±0.031 | 2.47±0.032 | 0.0025 |
| E, mm/s | 617±15.4 | 525±11.6 | <0.0001 |
| A, mm/s | 413±11.7 | 365±9.6 | 0.0056 |
| E slope | -36698±1832.1 | -32172±968.7 | 0.022 |
| E/A | 1.50±0.032 | 1.46±0.039 | 0.59 |
| IVRT, ms | 16.1±0.40 | 16.1±0.22 | 0.70 |
| IVCT, ms | 11.6±0.30 | 12.1±0.18 | 0.70 |
| MPI | 0.69±0.016 | 0.72±0.013 | 0.20 |
| E', mm/s | -29.0±0.69 | -23.7±0.57 | <0.0001 |
| A', mm/s | -17.9±0.52 | -15.5±0.46 | 0.0030 |
| E'/A' | 1.63±0.034 | 1.56±0.046 | 0.40 |
| E/E' | -21.4±0.69 | -22.5±0.58 | 0.29 |

### The extent of left ventricle reverse remodelling after cessation of the MHS depends on the diet in female mice

We previously showed that stopping AngII, shifting to a low-fat diet (LFD) and introducing voluntary exercise (VE) led to significant LV reverse remodelling after the MHS in both male and female mice (4). This lifestyle correction also helped reverse these mice’s exercise capacity loss. Here, we compared the effects of stopping AngII, the primary hypertrophic stress, without introducing VE and feeding the mice with the LFD (Sedentary RR or Sed group) or continuing the HFD while introducing VE (Trained HFD or HFD group). These mice were compared to control (Ctrl group) mice (28 days sedentary and fed the standard diet, then 28 days with VE and LFD) and MHS mice (28 days of AngII and HFD, then 28 days with VE and LFD) (RR group) (Figure 1A). Figure 1B illustrates representative long-axis heart sections stained with picrosirius red of females (top) and males (bottom) from the different groups. As previously observed, the MHS stress did not change body weight after 28 days but resulted in significant cardiac hypertrophy (Figure 1C). Stopping the HFD kept body weight at the level of MHS mice at the end of 28 days (Figure S1). Mice which continued being fed the HFD were heavier (Figure S1). After 28 days, MHS significantly increased heart weight (indexed for tibial length; iHeart) in both males and females (Figure 1C and dotted lines in Figure 1D). Normalization of indexed heart weight (iHeart) was observed in all three recovery groups (RR, Sed and HFD) (Figure 1D and Figure S1). In HFD female mice, the indexed heart weight (iHeart) was not normalized compared to control animals (Ctrl) and remained higher than the RR and Sed mice. As expected and previously observed, left atrial (LA) enlargement was present in male and female mice after MHS (Figures 1E, dotted lines). Four weeks of recovery did not normalize LA weight in males but did in females. Sed and HFD male animals still had a small degree of LA enlargement compared to controls. In females, changing the diet to LFD was sufficient to reduce LA weight to control levels, but continuing HFD blocked normalization. We measured blood pressure (BP) in conscious mice, as illustrated in Figure 1F. The MHS raised mean BP in female and male animals, and 28 days after stopping AngII, the BP returned to normal except for females in the HFD group. BP in females was higher than in males.

### Reverse LV remodelling is associated with a loss of ejection fraction in male mice

After 28 days of recovery following the MHS, we studied LV remodelling and function by Echo. As illustrated in Figure 1F, LV wall thickness (septal wall + posterior wall thickness) in all male and female groups was like control animals except for HFD males, which remained as measured immediately after the MHS. A tendency for thicker LV walls in RR, Sed and HFD compared to controls was observed, however (p=0.073, p=0.092 and p=0.092, respectively) LV ejection fraction (EF) remained unchanged after MHS. EF remained unchanged in all female groups after recovery. On the other hand, EF values in male RR and Sed groups were decreased compared to controls and MHS (Figure 1G). LV’s internal dimensions changed during recovery in a sex-specific manner. In males, end-diastolic diameter (EDD) was unchanged by MHS and remained like controls after four weeks of recovery (Figures 1H and 1I). End-systolic diameter (ESD) was increased in the RR and Sed groups, explaining EF losses (Figure 1I). EDD and ESD values were increased after recovery compared to the MHS group in females (Figures 1GH and 1I).

Figure S2 shows that both E and A waves remained unchanged in male and female animals after recovery, except for the RR groups, which displayed higher E and A values. The E/A ratio and intra-ventricular relaxing time showed no change (Figure S2). More complete echocardiography data are in Tables S2 and S3.

Cardiomyocyte cross-sectional area (CSA) did not return to normal in males 28 days after the MHS. However, all recovery groups showed a reduction compared to MHS CSA values but not normalization (Figure S3A). In females, stopping AngII and the HFD (Sed group) returned CSA to control levels. Only the Sed group showed normal CSA levels, suggesting VE could help maintain cardiomyocyte hypertrophy. Continuing HFD for four additional weeks kept cardiomyocyte size at MHS levels. Myocardial interstitial fibrosis levels remained unchanged in females after recovery, whereas they were slightly reduced in males’ RR and Sed groups (Figure S3A). LV sections stained with WGA for CSA measurements and picrosirius red for fibrosis are also illustrated (Figures S3B and S3C).

### Cardiac hypertrophy and extracellular matrix (ECM) remodelling markers gene expression are not sensitive tools for assessing the extent of myocardial recovery

We assayed the expression levels of several LV hypertrophy or fibrosis marker genes to determine if they correlated with the reverse remodelling observed in the mice. As illustrated in Figures 2A and B, LV gene expression of natriuretic peptides, *Nppa* (atrial) and *Nppb* (brain), was increased by MHS. This increase was more substantial for *Nppa* in females and *Nppb* in males, showing a sex dimorphism. Only in HFD groups were *Nppa* levels normalized, whereas Nppb mRNA levels for all recovery groups (RR, Sed and HFD) were like those of the control group. Connexin 43 (*Gja1*) mRNA levels were normalized in all recovery groups (RR, Sed and HFD) (Figure 2C).

**Figure 2.**
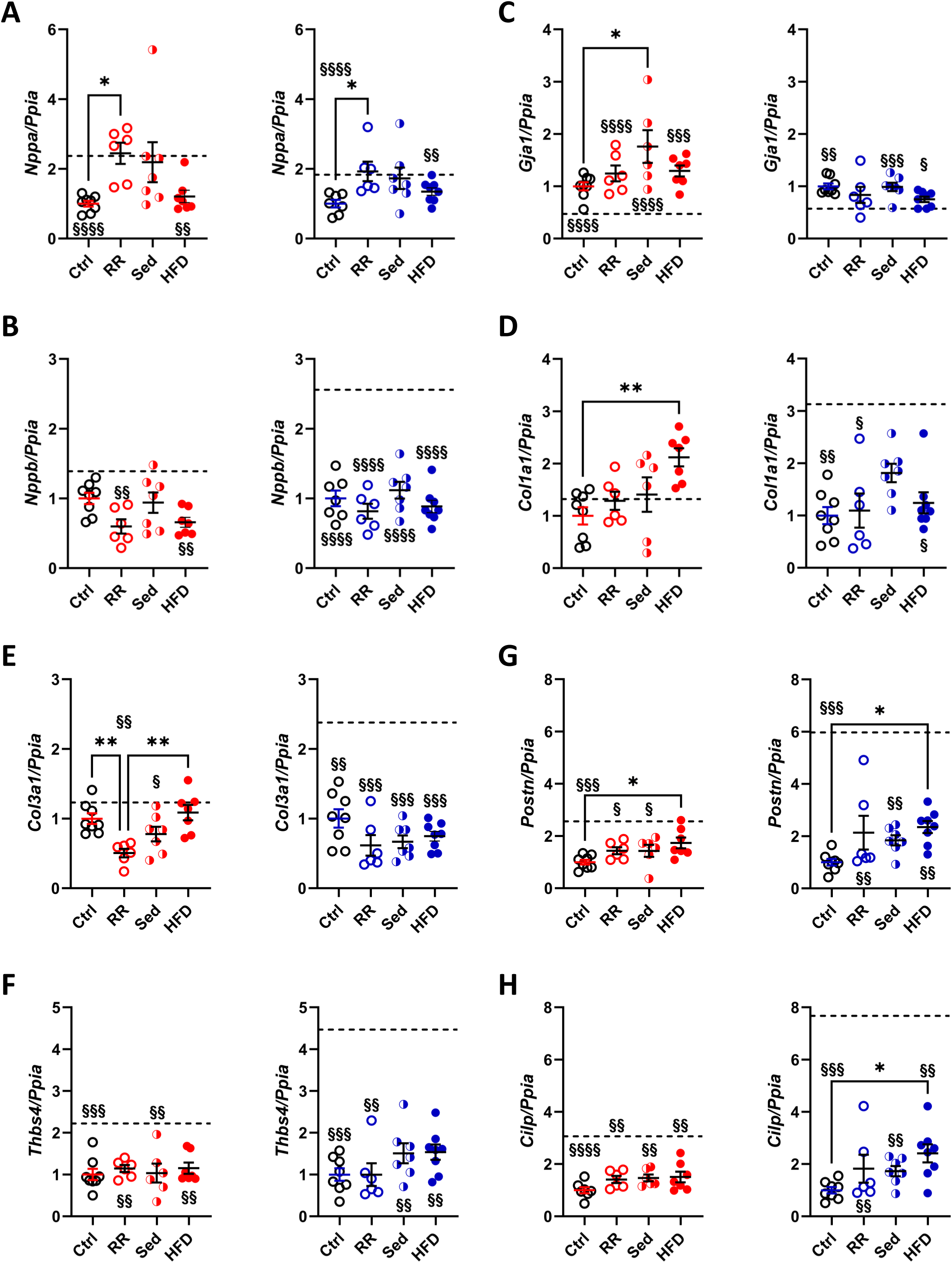
Effects of three myocardial recovery regimens after MHS Ctrl: control, RR: reverse remodelling group, Sed: LFD without voluntary exercise (VE) and HFD: HFD and VE on expression levels of LV hypertrophy and myocardial fibrosis genes. The dotted line on a graph represents the value of the indicated parameter after 28 days of MHS. The graph on the left is for females (red), and the one on the right is for males (blue). A. *Nppa*, atrial natriuretic peptide. B. *Nppb*, brain natriuretic peptide. C. *Gja1*, connexin 43, D. *Col1a1*, Collagen 1 α1, E. *Col3a1*, Collagen 3 α1, F. *Tbsp4*, thrombospondin 4. G. *Postn*, periostin and H. *Cilp*, cartilage intermediate layer protein. Data are represented as mean +SEM (n=5-6 per group). One-way ANOVA followed by Holm-Sidak post-test. *: p<0.05, **: p<0.01, ***: p<0.001 and ****: p<0.0001 between indicated groups. §: p<0.05, §§: p<0.01, §§§: p<0.001 and §§§§: p<0.0001 between the indicated group and MHS group using the Student’s T-test.

Procollagens 1 and 3 genes (*Col1a1* and *Col3a1*) were increased in males after MHS and normalized in the recovery groups (Figures 2D and 2E). In females, however, MHS did not significantly increase the expression levels of these two genes. *Col1a1* was increased in the female HFD group, and Col3a1 mRNA levels were below the controls of the RR and Sed groups. Thrombospondin 4 (*Thbs4*), Periostin (*Postn*) and Cartilage intermediate layer protein 1 (*Cilp*), marker genes of extracellular matrix remodelling, had their expression levels significantly increased by MHS and normalized in all recovery groups except for *Postn* in HFD females (Figures 2G and 2H). Interestingly, expression levels of all ECM markers increased after MHS in males more than in females (Figures 2E to 2H).

### The female heart is more resilient to a second MHS

As schematized in Figure 3A, we exposed male and female animals to a second MHS after an extended recovery period of 12 weeks instead of four. We hoped to evaluate the extent of myocardial recovery after 12 weeks and the cardiac resiliency to exposure to a second round of the AngII/HFD regimen (MHS) afterward. Controls were fed a standard diet (17% calories from fat) and maintained sedentary from day 1 to 28. Then, for 12 weeks, they were fed the low-fat diet (LFD; 10% fat) and allowed to VE from day 29 until day 140 to match the conditions of the 12-week recovery group (MR). During the 12 weeks of recovery following the first MHS, the MR group mice were thus fed LFD and could VE. Half the mice were euthanized, and the other half were exposed to a second MHS for 28 days (MR + MHS). Figure 3B illustrates representative long-axis LV sections stained with picrosirius red of males and females from these different groups.

**Figure 3.**
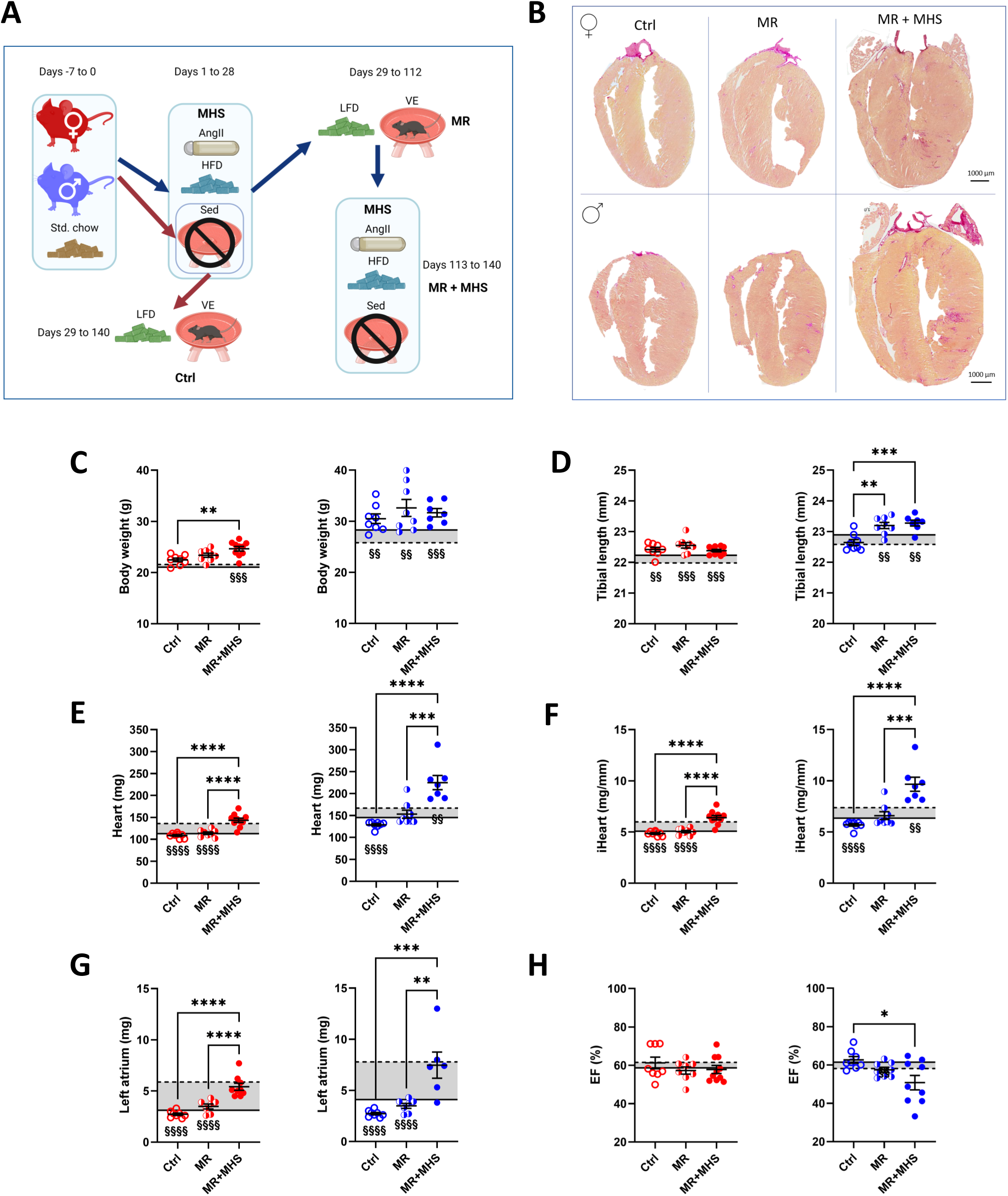
The male mouse’s heart is less resilient to a second MHS stress after 12 weeks of recovery. A. Longer myocardial recovery (**MR**) and second MHS. Male and female mice (8-week-old) were divided into Controls (**Ctrl**), **MR**, and **MR + MHS**. Controls: Mice were kept sedentary until they were 12 weeks old, then the animals were allowed voluntary exercise (VE) for 12 weeks and fed a low-fat diet (LFD) (red arrows). MR: MHS for 28 days, then RR for 12 weeks. MR + MHS: After RR for 12 weeks, a second 28-day MHS was performed (blue arrows). Created in BioRender. Couet, J. (2025) https://BioRender.com/o64o067)B. Representative images of picrosirius red staining of female and male heart sections for each indicated group. Effects of a longer recovery period (RR; 12 vs. 4 weeks) on the body weight (C), the tibial length (D), the heart weight (E), indexed heart weight (F; iHeart) and left atrial weight (G) and the LV ejection fraction (H). The solid line on a graph represents the value of the indicated parameter in mice in 12-week-old control mice, and the dotted line is after the first MHS. One-way ANOVA followed by Holm-Sidak post-test. *: p<0.05, **: p<0.01, ***: p<0.001 and ****: p<0.0001 between indicated groups. §: p<0.05, §§: p<0.01, §§§: p<0.001 and §§§§: p<0.0001 between the indicated group and MHS group using the Student’s T-test.

As illustrated in Figures 3C and 3D, female and male mice were larger and heavier after 12 weeks of recovery than younger mice (shaded area on the graphs), as expected. The extended period of recovery (12 weeks instead of four) reversed the cardiac hypertrophy caused by the MHS in females, whereas in males, this reversal was less complete (p=0.1) (Figures 3E and 3F). The second MHS caused similar levels of cardiac hypertrophy in females as for the first stress 12 weeks before (dotted line). In males, hypertrophy levels surpassed those recorded after the first MHS (dotted line). As mentioned above, three males died during the second MHS. Left atrial enlargement after the second MHS were relatively similar to those registered after the first stress (Figure 3G). However, males’ LV ejection fraction levels significantly decreased after the second MHS (Figure 3H).

Twelve weeks of recovery helped normalize LV wall thickness in male and female animals (Figure 4A), although they tended to be thicker than controls (p=0.069) in males. LV diameters (diastolic and systolic) were longer in MR males than in controls (Figures 4B and C). Those of MR females were normalized. The second MHS stress (MR+MHS) resulted in the thickening of the LV walls in males. In females, thickness was left unchanged (Figure 4A) after the second MHS. End-diastolic volume tended to be larger in MR males than in controls (P=0.055) and increased significantly after the second MHS. In females, however, LV volume remained stable (Figure 4D). Stroke volume and cardiac output were maintained in 12-week-recovery groups (male and female) after MHS (Table S4).

**Figure 4.**
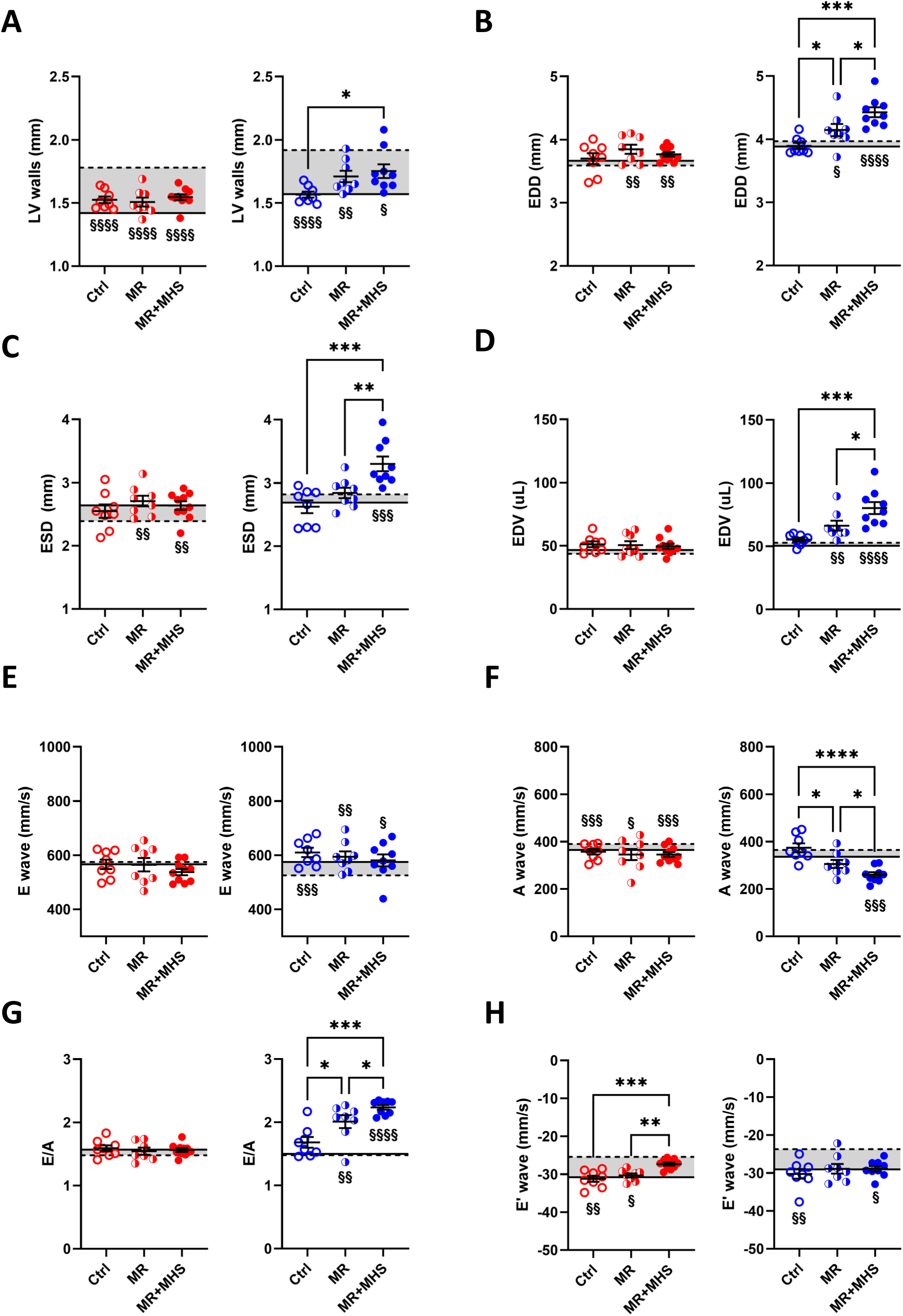
Echocardiography data of mice after 12 weeks of recovery (MR) and the second MHS (MR+MHS). The solid line on a graph represents the value of the indicated parameter in mice in 12-week-old control mice, and the dotted line is after the first MHS. A. LV wall thickness in diastole (PWd+IVSd) of female (red) and male (blue) mice. B. End-diastolic LV diameter (EDD). C. End-systolic LV diameter (ESD) and D. End-diastolic LV volume (EDV). E. E wave, F. A wave, G. E/A and H. E’ wave. One-way ANOVA followed by Holm-Sidak post-test. *: p<0.05, **: p<0.01, ***: p<0.001 and ****: p<0.0001 between indicated groups. §: p<0.05, §§: p<0.01, §§§: p<0.001 and §§§§: p<0.0001 between the indicated group and MHS group using the Student’s T-test.

Diastolic parameters in females were the same between the MR and MR+MHS groups and controls except for the E’ wave after the second stress (Figure 4E-F). In males, however, the A wave and the E/A ratio did not return to normal after 12 weeks and worsened even more after the second MHS.

The second MHS resulted in even more increased cardiomyocyte hypertrophy and myocardial interstitial fibrosis than the first stress, as illustrated in Figures 5A and B. LV sections labelled with fluorescent wheat germ agglutinin (WGA) show that the second MHS after 12 weeks of myocardial recovery resulted in a significant increase in cardiomyocytes CSA (Figure 5C). Staining was generally more abundant on the posterior LV wall than on the interventricular septum. Interestingly, the total amount of fibrosis staining was normalized after 12 weeks of recovery (MR) in males, whereas it remained unchanged in females. Twelve weeks of VE in normal mice (Ctrl) was associated with increased picrosirius red staining compared to young controls (solid line). The dotted line represents CSA or fibrosis after MHS.

**Figure 5.**
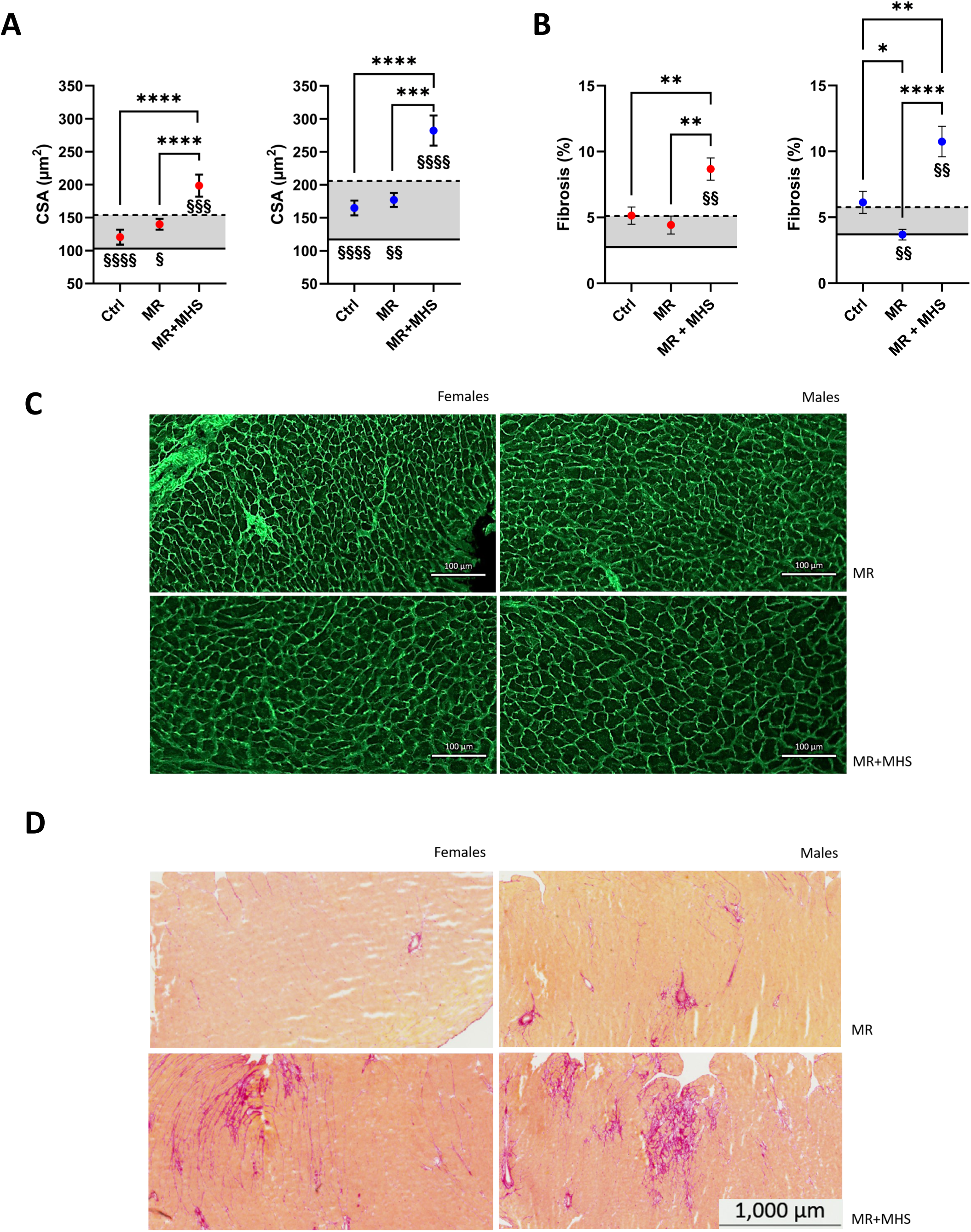
The second MHS results in additional cardiomyocyte hypertrophy and myocardial fibrosis. Females (red) and males (blue). A. Cross-sectional area of cardiomyocytes quantified by WGA-FITC staining (20-30 cells per mouse and six animals per group). The solid line on a graph represents the value of the indicated parameter in mice in 12-week-old control mice, and the dotted line is after the first MHS. Control animals after 12 weeks of VE. MR: MHS animals after 12 weeks of recovery and MR + MHS: mice after 2^nd^ MHS. B. Myocardial fibrosis (picrosirius red staining). C. Representative images of WGA-FITC staining of male and female MR and MR + MHS. D. Representative images of picrosirius red staining of female and male heart sections for each indicated group. One-way ANOVA followed by Holm-Sidak post-test. *: p<0.05, **: p<0.01, ***: p<0.001 and ****: p<0.0001 between indicated groups. §: p<0.05, §§: p<0.01, §§§: p<0.001 and §§§§: p<0.0001 between the indicated group and MHS group using the Student’s T-test.

We tested the same hypertrophy and ECM remodelling marker genes in mice after the second MHS. For the *Nppa, Col1a1, Col3a, Thbs4, Postn, and Cilp genes,* the second MHS more modulated mRNA levels than the first in female and male mice (Figure 6).

**Figure 6.**
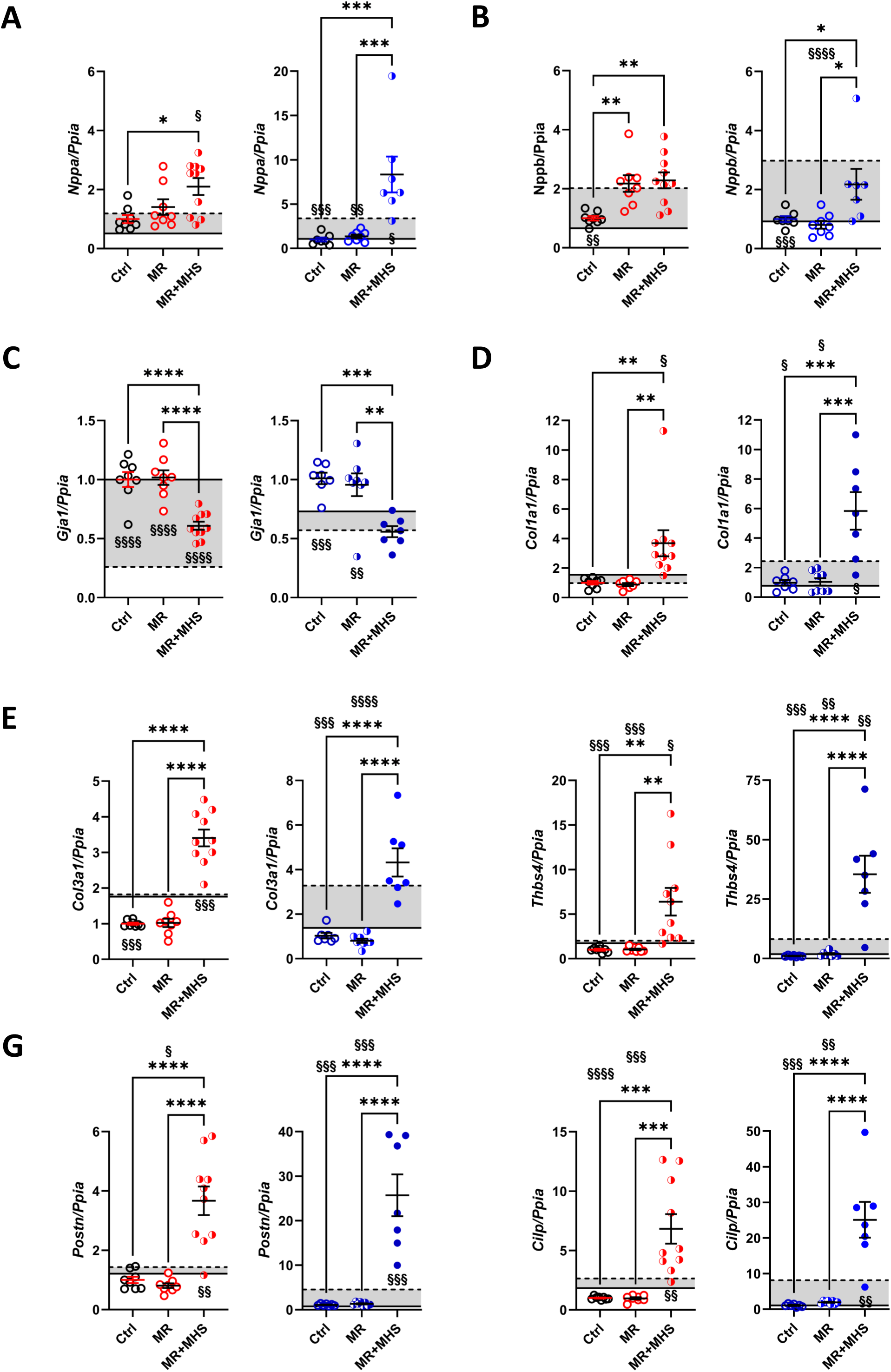
The second MHS results in a larger modulation of cardiac hypertrophy and fibrosis gene markers. The solid line on a graph represents the value of the indicated parameter in mice in 12-week-old control mice, and the dotted line is after the first MHS. A. *Nppa*, atrial natriuretic peptide. B. *Nppb*, brain natriuretic peptide. C. *Gja1*, connexin 43, D. *Col1a1*, Collagen 1 α1, E. *Col3a1*, Collagen 3 α1, F. *Tbsp4*, thrombospondin 4. G. *Postn*, periostin and H. *Cilp*, cartilage intermediate layer protein. Data are represented as mean +SEM (n=5-6 per group). One-way ANOVA followed by Holm-Sidak post-test. *: p<0.05, **: p<0.01, ***: p<0.001 and ****: p<0.0001 between indicated groups.

### Voluntary exercise (VE) for 12 weeks and a low-fat diet result in smaller hearts

As mentioned, the control mice were allowed to VE for 12 weeks and fed an LFD (10% calories from fat) from 3 months of age. We compared those animals to sedentary mice of the same age on a standard diet (17% calories from fat) to evaluate the impact of training and the LFD on the hearts of otherwise normal animals. As illustrated in Figure S4, VE did not influence males’ body weight or tibial length. Trained female mice were more petite than sedentary females. Heart weight was lower in trained animals of both sexes. Using echo, we observed that VE animals had smaller LV diameters and volumes than sedentary ones, resulting in concentric remodelling and decreased mass (Table S5). VE increased ejection fraction and global circumferential strain (GCS) in female mice (Table S5).

Diastolic parameters adaptations to VE and LFD were mild, as seen in Table S5. In males, VE resulted in lower A wave, shorter intra-ventricular contraction times (IVCT), and higher E’/A’ ratios. The left atrial diameter was reduced, and both tissue Doppler E’ and A’ waves were modulated by training, resulting in an increased E’/A’ ratio in females (Table S6).

### The MHS strongly increases the pyruvate dehydrogenase kinase 4 (PDK4) protein content in HFpEF mice

Recent observations in a different murine HFpEF model showed that myocardial substrate choice for energy production dramatically shifts toward fatty acids utilization (17). We were interested to see if this shift in myocardial energy substrate preference was present in the MHS HFpEF model and if the myocardial recovery period could help normalize it.

We first measured mRNA levels of genes encoding for fatty acids (*CD36/FAT*) and glucose entry (*Glut4* and *Glut1*) into the cell. The MHS resulted in lower mRNA levels for *Glut4* but not *CD36/FAT* or *Glut1* (Figures 7A to C). Myocardial recovery helped normalize *Glut4* expression levels. The second MHS again lowered *Glut4* levels in both sexes and raised *CD36/FAT* in females. Interestingly, Glut1 levels seemed to be increased by VE and the LFD in both male and female mice (Figure 7C). *Pdk4* gene expression (Figure 7D) was increased after MHS in male and female mice. Twelve weeks of recovery normalized *Pdk4* mRNA levels. The second MHS also decreased *Glut4* mRNA levels and did the opposite for those of *Pdk4*.

**Figure 7.**
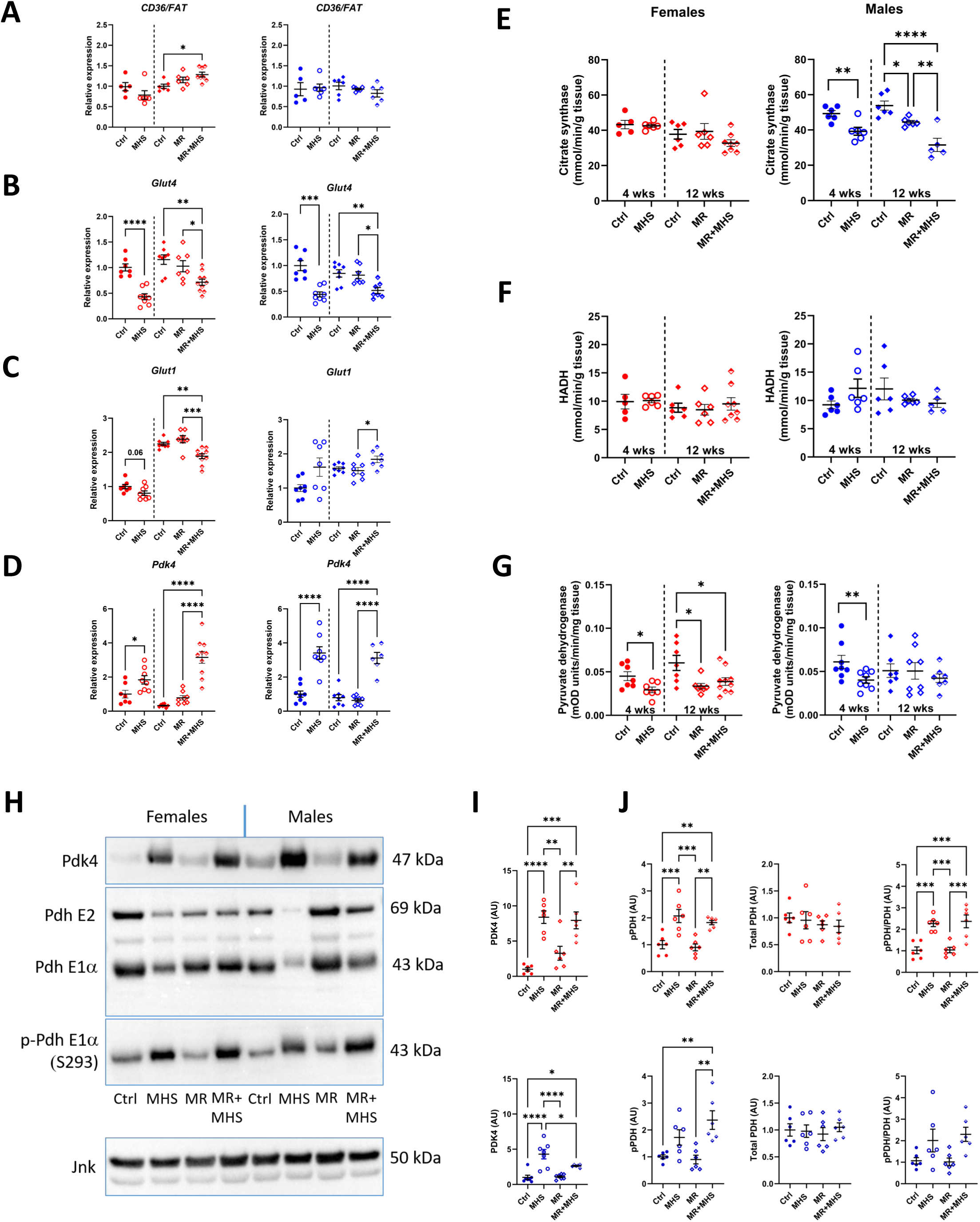
The MHS inhibits myocardial energy metabolism via the inhibition of glucose utilization. A-D. Expression levels of genes implicated in myocardial energy production. A. *CD36/FAT, fatty acid transporter*. B. *Glut4*, glucose transporter 4. C. *Glut1*, glucose transporter 1 D. *Pdk4*, pyruvate dehydrogenase kinase 4. Left, Ctrl: young control mice, MHS: Initial MHS stress. Student T-test. *: p<0.05, **: p<0.01, ***: p<0.001 and ****: p<0.0001 between indicated groups. Right: Ctrl: control mice after 12 weeks of training, MR: MHS mice after 12 weeks of recovery and MR + MHS: mice after 2^nd^ MHS. One-way ANOVA followed by Holm-Sidak post-test. *: p<0.05, **: p<0.01, ***: p<0.001 and ****: p<0.0001 between indicated groups. E, Citrate synthase activity, F, HADH activity and G, Pyruvate dehydrogenase activity. H. Representative blots of Pdk4, PDH, p-PDH and JNK content in young controls (Ctrl), MHS, MR and MR + MHS female (left) and male (right) mice. I. Quantification of Pdk4 content. J. Quantification of p-PHD, total PDH and p-PDH/PDH contents.

Figure 7E illustrates that myocardial citrate synthase activity, the first step of the citric acid cycle and often a marker of mitochondrial function, was reduced by the MHS in males but not in females. Twelve weeks of recovery were insufficient to return citrate synthase to normal levels, and they were further reduced after the second MHS. This was not observed in females. Hydroxyacyl-Coenzyme A dehydrogenase activity, implicated in fatty acid beta-oxidation, was not modulated by the MHS stress or during recovery (Figure 7F).

The pyruvate dehydrogenase E1 subunit (PDH E1) is part of the pyruvate dehydrogenase complex and catalyzes the conversion of pyruvate into acetyl-CoA. The MHS reduced PDH myocardial activity in both female and male mice (Figure 7G). Twelve weeks of recovery were insufficient to normalize female activity levels, which remained low after the second MHS. This was not observed in males.

PDK4 phosphorylates the PDH E1 subunit, inhibiting its activity. As illustrated in Figures 7H and 7I, either the first or second MHS stress increased PDK4 levels significantly, resulting in higher levels of phosphorylation of the PDH E1 subunit (Figures 7H and 7J).

We then measured the gene expression levels of *CD36/FAT, Glut1, Glut4*, and *Pdk4* after four weeks of recovery and the effects of exercise and HFD on recovery. As illustrated in Figure S5, glucose transporters mRNA levels were normalized in all recovery groups, even over controls in females. *Pdk4* mRNA levels and protein content were not normalized in mice fed the HFD, and *CD36/FAT* mRNA levels were more elevated in mice fed the HFD.

## Discussion

Using a previously described HFpEF two-hit mouse model combining AngII continuous infusion and an HFD for 28 days (MHS) (4), we studied factors such as diet or physical activity that could influence cardiac RR once the hypertensive stress (AngII) stopped. We showed in male mice that stopping AngII was the main factor influencing RR at the morphological level, irrespective of the diet or the VE. In females, continuing the HFD during recovery maintained cardiac hypertrophy. An extended recovery period (12 vs. four weeks) led to a seemingly more complete RR in females but not males. Males were thus more sensitive to a second MHS than females after 12 weeks of recovery and showed a possible evolution toward HFrEF with LV dilation and loss of ejection fraction.

We had previously observed extensive reverse remodelling when the MHS was stopped, and a low-fat diet and voluntary exercise were introduced (4). Myocardial recovery was still incomplete, as some left atrial enlargement was present after four weeks. Exercise tolerance, although greatly improved, remained inferior to control animals. Here, we want to study independently the role of VE and HFD on myocardial recovery after stopping the primary hypertrophic stress, AngII. Our present results emphasize the complex interplay between these factors on myocardial recovery and that usual gene expression markers of hypertrophy and remodelling failed at identifying remaining functional or morphological anomalies, as evidenced by their normalization in both female and male mice after 12 weeks of recovery. In addition, sex differences were present during cardiac RR, and again, usual gene expression markers did not reflect macroscopic morphological and functional differences. For instance, LV collagens mRNA levels did not correlate with increased myocardial fibrosis in females.

We observed that cardiac hypertrophy and left atrial enlargement induced by MHS were reversed after four weeks of RR. This was also true for most LV marker genes analyzed, except for the atrial natriuretic peptide gene (*Nppa*). Myocardial fibrosis and cardiomyocyte surface area both remained increased compared to controls. We had previously shown that the LV exome was largely normalized after four weeks (4) but not the myocardial microRNA profile, which remained mostly unchanged (18). This emphasizes that the LV gene expression return to basal levels was not necessarily indicative of normalization but mainly a reflection of the causal stress cessation. Four weeks of recovery were possibly insufficient, or some abnormalities caused by the MHS may have become permanent. To answer this, we prolonged the recovery duration to 12 weeks and then tested its extent by administering a second MHS.

For most morphological and functional parameters tested after 12 weeks of recovery, female mice showed normalization compared to controls. However, in males, a tendency for a larger heart, thicker LV walls, and increased LV volume was observed. Although left atrial enlargement was reversed in males, the A wave, as measured using a power Doppler echo, was lowered after 12 weeks of recovery, resulting in an increased E/A ratio over 2, indicating a possible fall in atrial contractility. For all LV gene markers tested, expression levels were normalized entirely after 12 weeks of VE and LFD.

Our study highlighted several sex differences related to the response to the MHS and the myocardial recovery afterwards. For instance, the expression levels measured in six of the eight genes studied were more modulated in males than in females (*Nppb, Thbs4, Col1, Col3, Postn* and *Cilp*). It is difficult to estimate if the MHS has the same severity for both sexes. Increases in cardiac hypertrophy, cardiomyocyte hypertrophy and fibrosis are equivalent. LV concentric remodelling is similar. Diastolic function changes caused by the MHS are also identical. The recovery helped uncover sex differences, where males first lost the ejection fraction but recovered with time. The significance of this increased modulation of expression levels is still unclear. It may indicate that even though the recovery period was generally sufficient to normalize mRNA levels, they may have taken more time to return to basic levels. Interestingly, after 28 days, the MHS did not induce collagen 1 and 3 expression in females but did in males. It is possible that the collagen expression was transiently modulated in females during the MHS, resulting in increased myocardial fibrosis or slowing their degradation.

It is considered that male and female rodents adapt differently to a pathological stimulus. For instance, males exposed to a pressure overload often had more cardiac hypertrophy and fibrosis and evolved more rapidly toward heart failure and mortality than females (18–24). This is also true for other pathological stimuli, such as in models of hypertension, such as the SHR rat (25–28), the Dahl Salt-Sensitive rat (29–32), or myocardial infarction (33–40).

More recently, in murine multi-morbidity HFpEF models, sex differences were described in L-NAME+HFD-treated C57Bl6/N mice (41, 42), with females being resistant to the development of HF. However, this resistance was not observed in C57Bl6/J females (43).

In a multi-hit mouse model, combining AngII, HFD and old age, Withaar and Coll. showed that females developed HFpEF, but males evolved toward HFrEF instead (44). A similar observation was made in aging mice fed or not with a Western diet. Females on the diet maintained their ejection fraction compared to age-matched control but not males (45). In another aging HFpEF model, in rats carrying the mouse Ren-2 renin gene, higher mortality was observed in male rats and a significant loss of EF in surviving animals (46).

In another mouse model, this time in db/db and receiving aldosterone compared to wild-type mice, it was observed that this model recapitulates sex-specific features in HFpEF, worse diastolic dysfunction in females, and higher arrhythmia susceptibility in males (47). Also, in db/db mice, sex differences in the development of the HFpEF at 6 months were observed. Female animals had increased weight gain and higher levels of hypertension, with worse diastolic dysfunction. An expansion of the perivascular collagen network was also described. In males, however, microvascular rarefaction was observed (48).

The ejection fraction slightly decreased after recovery in males as their LV diameter and volume increased during recovery, and LV walls became thinner. Importantly, cardiac output, as evaluated by echocardiography, was maintained for both male and female animals. Still, the morphological changes during LV reverse remodelling differed between the sexes and showed a potential evolution in males toward LV dilation. This transient loss in ejection fraction in males was absent when they were maintained on the HFD and had more concentric remodelling.

We had previously shown that the high-fat diet was hypertrophic for the female heart (4), and here, we observed that it slowed reverse remodelling only in females. The HFD increased gene markers of fibrosis, such as collagen 1, collagen 3 and periostin. Myocardial fibrosis levels were not corrected in females after four weeks of recovery when the HFD was maintained, and myocyte hypertrophy was not reduced either. In males, this effect on reverse remodelling of the HFD was not observed. This suggests that young females’ hearts are more responsive to the high-fat diet than males, especially for the hypertrophic response and the resulting myocardial fibrotic response.

The impact of voluntary exercise is more subtle in cardiac RR after the MHS. When both the AngII and the HFD stopped, it was unclear whether VE provided an additional advantage over the Sed during recovery. The main difference observed between active and inactive recovering mice was at the level of several echocardiography diastolic parameters. Active mice of both sexes had increased E wave and A wave values compared to inactive animals. We let mice voluntarily exercise after stopping AngII and HFD for either 4 or 12 weeks. Heart weight returned to control levels in females, whereas cardiac hypertrophy was only partially reversed in males. A more extended recovery period helped reverse left atrial enlargement to near normal. It is thus challenging to conclude that VE impacted our mice, although we showed that it had a clear and surprising impact in normal mice.

For the 12-week recovery period, our control animals also voluntarily exercised and were fed a low-fat diet (10% of calories from fat). Unexpectedly, we observed a marked heart response in these animals compared to age-matched inactive mice fed a standard diet (17% of calories from fat). We were expecting some levels of physiological cardiac hypertrophy in trained animals, especially females (49). Instead, trained animals’ heart weights were smaller, LV diameters shorter, LV volumes diminished, and ejection fraction improved, especially in females. LV wall thickness was similar between inactive and active mice, as was cardiac output. This shows that voluntary exercise markedly affects our animals’ hearts and that diet may play a role. The current study was not designed to study the effects of VE on the normal heart, as the trained normal mice were employed as controls for the MR group. Thus, the LFD could have blocked the expected hypertrophic response to training or running distances in HFpEF, which were insufficient to induce this response.

Targeting physical activity, nutritional changes, and treating other causes of HFpEF have been shown to improve patients’ quality of life (50). HFpEF patients are often old and find it less easy to enroll in structured exercise programs, so this treatment option has not received all the attention it deserves. (50–53). Along with sodium-glucose cotransporter type 2 (SGLT2) inhibitors and glucagon-like peptide 1 (GLP1) agonists (54, 55), which are becoming a first-line treatment, cardiorespiratory rehabilitation, diet changes and management of hypertension are all required for efficient management of HFpEF. In this two-hit HFpEF mouse model, we showed that treating hypertension and dietary changes can benefit cardiac health and promote myocardial recovery, although incomplete.

We also reproduced in our mice observations made in the L-NAME+HFD model that glucose utilization is reduced as a substrate for myocardial ATP production via downregulation of PDH activity (17). In addition, we showed that RR was associated with the downregulation of Pdk4 content and, consequently, lower levels of PDH phosphorylation in both females and males. Continuing the HFD during RR blocked this normalization of the Pdk4-PDH axis. In the absence of AngII during RR, mice developed some levels of obesity, but those did not result in additional cardiac hypertrophy. We observed that an LFD would result in a better reversal of hypertrophy, as observed previously by Güven et Coll. (56), which showed that an LFD blocked both obesity and HFpEF development in their mouse model Because of the lipolytic action of AngII (4, 57), our model does not develop obesity only a little raise in the body fat content in females. We recently showed that the circulatory microRNA content of mice treated with AngII alone, the HFD or a combination of both, as in here, were relatively similar (58). It is thus possible that the adipose tissue, the largest producer of microRNAs in the circulatory system, has a similar stress response to insults.

We also observed that male mice had reduced citrate synthase activity after MHS but not females. This decrease in activity has been described before in two different murine HFpEF models (59, 60) and a rat model (61). Our observation and others point toward mitochondrial damages that possibly persist after RR and could cause the males to be more sensitive to a second MHS. This will have to be more thoroughly examined in further studies to evaluate damage in mitochondrial function in our model and the mitochondrial capacity to repair during recovery.

## Study limitations

As mentioned above, it is difficult to affirm that both sexes were impacted similarly by the MHS, and this may play a role in the levels of myocardial recovery achieved once the stress is stopped.

We did not monitor the mice’s running activity since they were not housed individually. It is possible that control mice trained more intensely than MHS mice during recovery. However, differences between trained control mice and MR animals were relatively mild at the macroscopic level.

This study was conducted in the C57Bl6/J mouse strain, as are most studies in the field. Studies conducted in inbred strains potentially have a more limited translational value. It will be important to confirm in various strains or in outbred mice if observations made in HFpEF models can be generalized or are strain-specific (43, 62).

## Conclusion

In conclusion, we showed that reverse remodelling after the MHS was stopped depended on correcting the HFD to the LFD in female mice. Cardiomyocyte hypertrophy and myocardial fibrosis did not decrease after four weeks of recovery. Males’ recovery after an extended period of 12 weeks seems insufficient to protect their hearts against a second stress, as well as in females. We observed that the gene expression of a series of markers was inadequate in identifying those male mice that recovered less than females. A diastolic function parameter such as the A wave and incomplete reversal of cardiac hypertrophy were more indicative of the persistence of male anomalies.

## Grants

The work was supported by grants from the Canadian Institutes for Health Research PJT-1665850 (to J. Couet and M. Arsenault) and from the Fondation de l’Institut universitaire de cardiologie et de pneumologie de Québec.

