## Supplemental data (Tables and figures) for "Relative contributions of correcting the diet and voluntary exercise to myocardial recovery in a two-hit murine model of heart failure with preserved ejection fraction"

**Table S1**. Sequences of primers used in this study.

| Symbol | Description | Forward sequence  Reverse sequence |
| --- | --- | --- |
| Cilp | Cartilage Intermediate Layer Protein | 5'-GCT TGT GTC CTG ATG GCA AA-3'  5'-CGA CGT GCT TTC ATC TCA GG-3' |
| Col1a1 | Collagen Type I Alpha 1 Chain | 5'-CAT TGT GTA TGC AGC TGA CTT C-3'  5’CGC AAA CAC TCT ACA TGT CTA GG-3’ |
| Col3a1 | Collagen Type III Alpha 1 Chain | 5'-TCT CTA GAC TCA TAG GAC TGA CC-3'  5’ TTC TTC TCA CCC TTC TTC ATC C-3’ |
| Fat/Cd36 | Fatty Acid Translocase/CD36 | 5'-GAT CGG AAC TGT GGG CTC AT-3'  5’-CTT TGC CAC GTC ATC TGG G 3’ |
| Gja1 | Connexin-43 | 5'-ACG GCA AGG TGA AGA TGA GA-3'  5'-GAG AGA CAC CAA GGA CAC CA-3' |
| Glut1 | Glucose Transporter Type 1 | 5'-TCT CTG TCG GCC TCT TTG TT-3'  5'-CGC AGT ACA CAC CGA TGA TG-3' |
| Glut4 | Glucose Transporter Type 4 | 5'-CAG TAC TCC CTG CTC TCC TG-3'  5'-GCT CTC TCT CCA ACT TCC GT-3' |
| Nppa | Natriuretic Peptide B | 5'-CTC CTT GGC TGT TAT CTT CGG-3'  5'-GGG TAG GAT TGA CAG GAT TGG-3' |
| Nppb | Natriuretic Peptide A | 5'-AGG TGA CAC ATA TCT CAA GCT G-3'  5’-CTT CCT ACA ACA TCA GTG C-3’ |
| Pdk4 | Pyruvate Dehydrogenase Kinase 4 | 5'-TTC TTG AAG AGT TCG AAG AGC A-3'  5'-CAT CGC CAG AAT TAA ACC TCA C-3' |
| Ppia | Cyclophilin a | 5'-TTC ACC TTC CCA AAG ACC AC-3'  5'-CAA ACA CAA ACG GTT CCC AG-3' |
| Postn | Periostin | 5'-GCT TTC GAG AAA CTG CCA CG-3'  5'-ATG GTC TCA AAC ACG GCT CC-3' |
| Thbs4 | Thrombospondin 4 | 5'-GAT ACT GAC GGG GAT GGG AG-3'  5'-CGT CAC TGT CTT GGT TGG TG-3' |

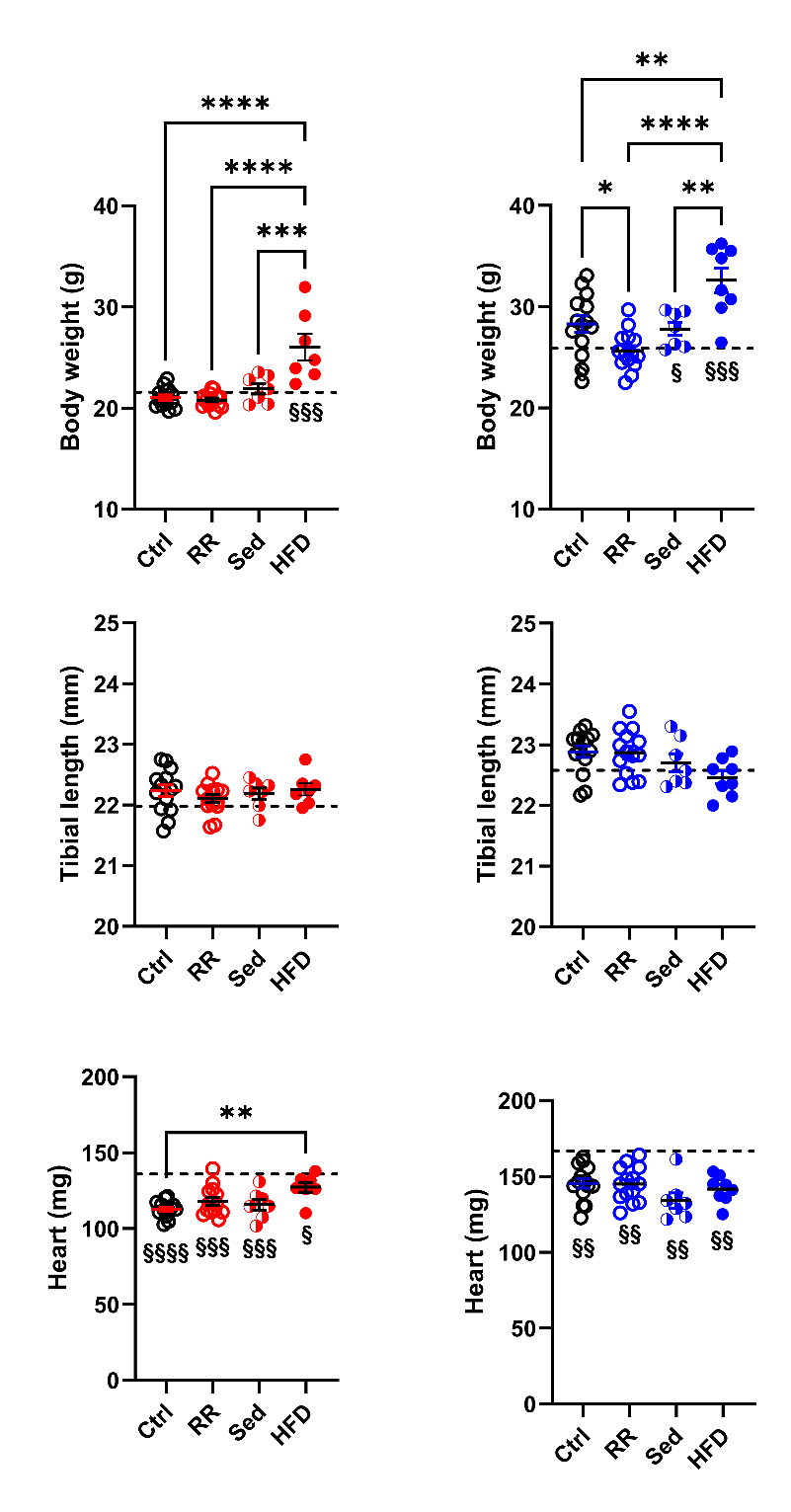

**Figure S1**. Body weight, tibial length and heart weight of mice after four weeks of recovery post-MHS. Females are red (left), and males are blue (right). Ctrl: control, RR: reverse remodelling group, Sed: LFD without voluntary exercise (VE) and HFD: HFD and VE. The dotted line represents the mean of the parameter after MHS and before recovery. One-way ANOVA followed by Holm-Sidak post-test. *: p<0.05, **: p<0.01, ***: p<0.001 and ****: p<0.0001 between indicated groups. §: p<0.05, §§: p<0.01, §§§: p<0.001 and §§§§: p<0.0001 between the indicated group and MHS group using the Student’s T-test.

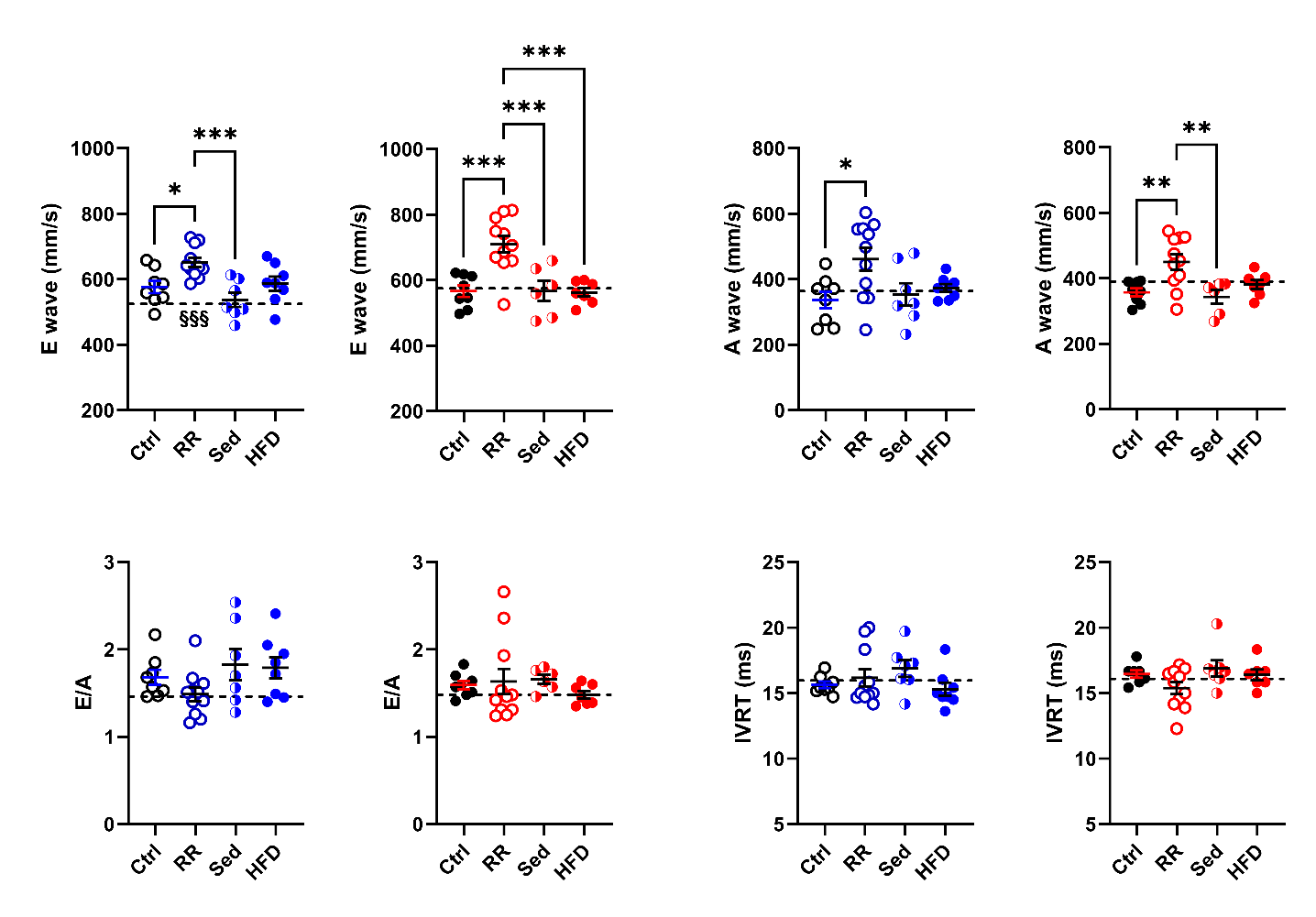

**Figure S2**. Echocardiography data of mice under three different regimens for myocardial recovery. Blue (left panels): Males and Red (right panels): Females. Ctrl: control, RR: reverse remodelling group, Sed: low-fat diet without voluntary exercise (VE) and HFD: HFD and VE. The dotted line on a graph represents the value of the indicated parameter after 28 days of MHS. IVRT: Isovolumetric relaxation time (IVRT). Data are represented as mean +SEM (n=7-8 per group). One-way ANOVA followed by Holm-Sidak post-test. *: p<0.05, **: p<0.01, ***: p<0.001 and ****: p<0.0001 between indicated groups. §§§: p<0.001 between the indicated and MHS groups using the Student’s T-test.

**Table S2.** Reverse remodelling (RR) after MHS: influence of correcting the diet and keeping the animals sedentary (Sed) or vice versa (HFD) on echo parameters. Echo data in male and female mice after MHS. Standard echo left ventricle parameters after four weeks of MHS and after four weeks of remission (RR, Sed or HFD). Control mice were studied in parallel Echo exams as described in the Methods section. PW: diastolic posterior wall thickness, IVS: diastolic inter-ventricular septum, RWT: relative wall thickness, EDV: end-diastolic volume, ESV: end-systolic volume, SV stroke volume, HR: heart rate, and CO: cardiac output. Results are expressed as the mean ± standard error of the mean (SEM). One-way ANOVA analysis and Holm-Sidak post-test. a: p<0.05, b: p<0.01, c: p<0.001 and d: p<0.0001 vs. control (Ctrl) group.

| **Males** |  |  |  |  |  |
| --- | --- | --- | --- | --- | --- |
| Parameters | Ctrl (n=8) | MHS (n=8) | RR (n=11) | Sed (n=7) | HFD (n=8) |
| PW, mm | 0.79±0.008 | 1.03±0.027^d^ | 0.83±0.028 | 0.83±0.016 | 0.95±0.022^d^ |
| IVS, mm | 0.78±0.008 | 0.91±0.023^d^ | 0.80±0.017 | 0.77±0.017 | 0.86±0.013^b^ |
| RWT | 0.41±0.008 | 0.53±0.019^d^ | 0.42±0.012 | 0.41±0.012 | 0.47±0.011^d^ |
| EDV, µl | 57±2.9 | 53±2.4 | 63±1.7 | 60±3.1 | 57±2.2 |
| ESV, µl | 22±1.2 | 22±1.6 | 31±1.3^d^ | 30±1.4^c^ | 23±1.5 |
| SV, mm | 35±2.1 | 31±1.4 | 32±0.8 | 30±2.4 | 34±1.3 |
| HR, bpm | 506±18.4 | 513±16.9 | 480±21.2 | 517±15.5 | 544±15.9 |
| CO, ml/min | 17.6±1.25 | 15.8±1.04 | 15.3±0.67 | 15.6±1.50 | 18.3±0.87 |
| **Females** |  |  |  |  |  |
| Parameters | Ctrl (n=7) | MHS (n=8) | RR (n=11) | Sed (n=7) | HFD (n=7) |
| PW, mm | 0.75±0.018 | 0.88±0.024^d^ | 0.82±0.026^c^ | 0.80±0.022 | 0.81±0.026 |
| IVS, mm | 0.67±0.022 | 0.84±0.015^d^ | 0.74±0.015^b^ | 0.75±0.012^b^ | 0.75±0.019^b^ |
| RWT | 0.39±0.014 | 0.48±0.011^d^ | 0.41±0.012 | 0.41±0.008 | 0.41±0.014 |
| EDV, µl | 42±1.6 | 47±1.2^a^ | 52±1.4^d^ | 46±3.1 | 45±2.3 |
| ESV, µl | 18±1.5 | 18±1.2 | 24±1.4^c^ | 21±1.6 | 18±1.4 |
| SV, mm | 25±1.0 | 30±2.4 | 28±1.0 | 25±3.2 | 27±1.2 |
| HR, bpm | 493±18.3 | 528±6.0^b^ | 505±16.5 | 527±42.7 | 513±13.8 |
| CO, ml/min | 12.3±0.85 | 15.7±0.84^a^ | 14.2±0.78 | 12.5±1.2 | 13.9±0.7 |

**Table S3.** Reverse remodelling (RR) after MHS: influence of correcting the diet and keeping the animals sedentary (Sed) or vice versa (HFD) on diastolic echo parameters. Echo data in male and female mice after MHS. Standard echo left ve’ntricle parameters after four weeks of MHS and after four weeks of remission (RR, Sed or HFD). E’: E wave and A’: A’ wave. Results are expressed as the mean ± standard error of the mean (SEM). One-way ANOVA analysis and Holm-Sidak post-test using data in young as controls. a: p<0.05, b: p<0.01, c: p<0.001 and d: p<0.0001 vs. control (Ctrl) group.

| **Males** |  |  |  |  |  |
| --- | --- | --- | --- | --- | --- |
| Parameters | Ctrl (n=8) | MHS (n=8) | RR (n=11) | Sed (n=7) | HFD (n=8) |
| E’ | -27.3±0.83 | -24.0±1.90 | -27.7±0.82 | -25.7±1.38 | -30.2±0.74 |
| A’ | -16.1±0.94 | -16.5±1.29 | -18.7±1.10 | -14.5±0.91 | -17.4±1.23 |
| E’/E | -21.3±0.96 | -23.1±1.51 | -23.7±0.54 | -21.2±1.11 | -19.4±0.73 |
| **Females** |  |  |  |  |  |
| Parameters | Ctrl (n=7) | MHS (n=8) | RR (n=11) | Sed (n=7) | HFD (n=7) |
| E’ | -30.0±1.10 | --27.0±1.14 | -30.1±1.10 | -27.2±1.44 | -27.1±0.82 |
| A’ | -16.4±0.64 | -17.7±0.98 | -21.0±0.89c | -16.9±1.16 | -16.8±0.88 |
| E’/E | -20.4±1.15 | -22.6±1.11 | -23.8±0.66 | -19.2±1.32 | 20.4±0.66 |

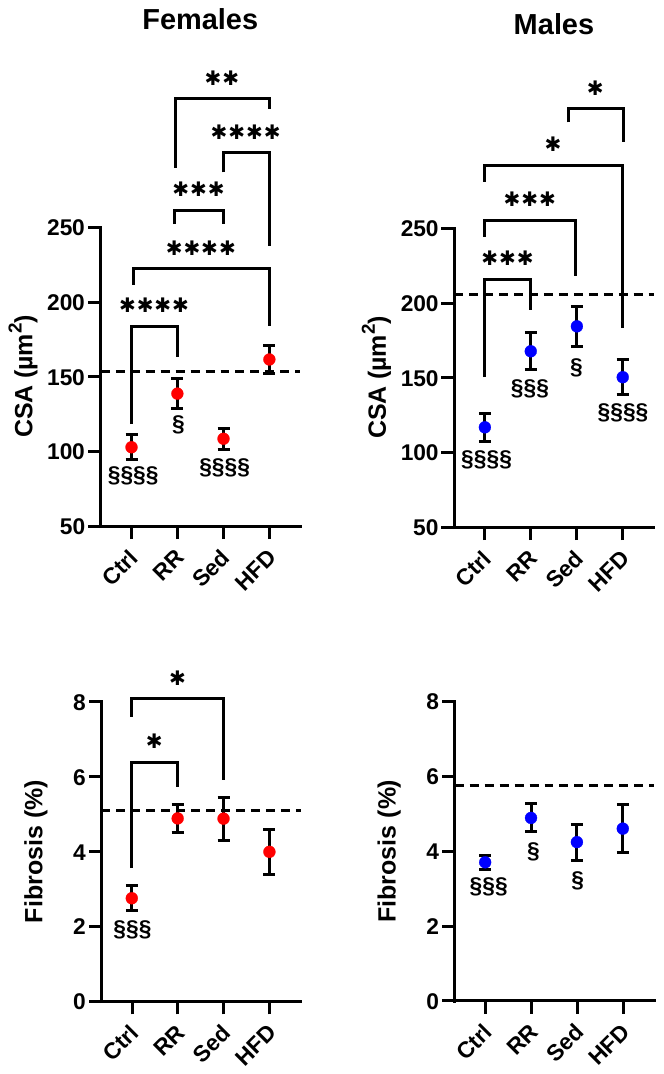

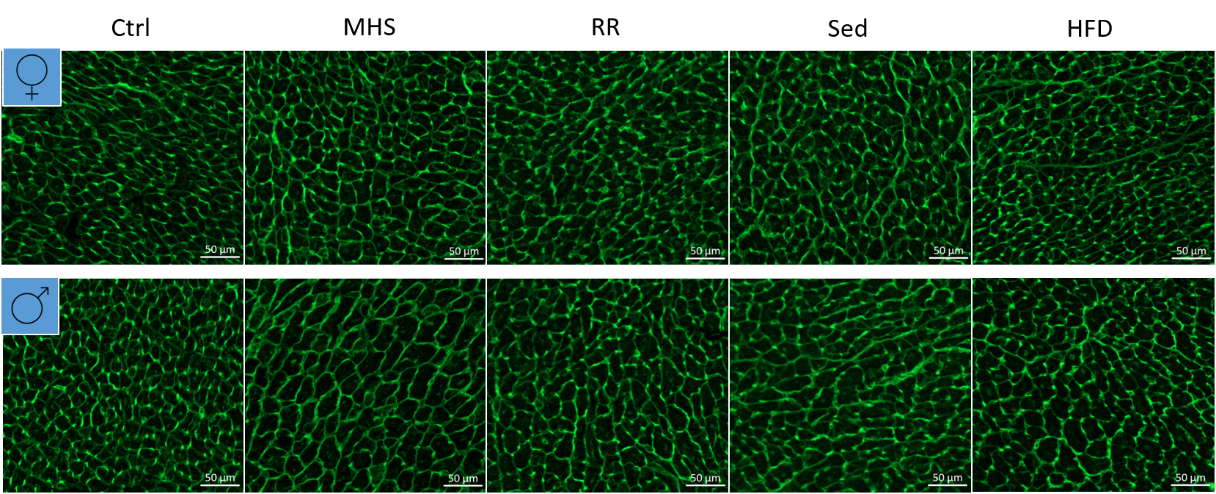

**Figure S3.** Reverse remodelling (RR) after MHS: influence of correcting the diet and keeping the animals sedentary (Sed) or vice versa (HFD) on cardiomyocytes cross-sectional area and myocardial fibrosis. Dotted line: mean of the parameter after 28 days of MHS. One-way ANOVA analysis and Holm-Sidak post-test using data in young as controls. a: p<0.05, b: p<0.01, c: p<0.001 and d: p<0.0001 vs. control (Ctrl) group. Representative images of WGA-FITC staining of male and female LV sections.

**Table S4.** Echocardiography parameters after a longer recovery (12 weeks) and a second MHS. During the twelve weeks of recovery following the first MHS, the mice were fed a low-fat diet (LFD) and could voluntarily exercise. Half the mice were euthanized (MR), and the other half were exposed to 28 days of MHS (MR + MHS). Control (Ctrl) animals were not exposed to the first MHS but kept sedentary for 4 weeks, and their standard diet was changed to the LFD for the following 12 weeks with VE Echo data in male and female mice after MHS. Standard echo left ventricle parameters after four weeks of MHS and after four weeks of remission (RR, Sed or HFD). Control mice were studied in parallel Echo exams as described in the Methods section. PW: diastolic posterior wall thickness, IVS: diastolic inter-ventricular septum, RWT: relative wall thickness, EDV: end-diastolic volume, ESV: end-systolic volume, SV stroke volume, HR: heart rate, and CO: cardiac output. Results are expressed as the mean ± standard error of the mean (SEM). One-way ANOVA analysis and Holm-Sidak post-test. a: p<0.05, b: p<0.01, c: p<0.001 and d: p<0.0001 vs. control (Ctrl) group.

| **Males** |  |  |  |
| --- | --- | --- | --- |
| Parameters | Ctrl (n=8) | MR (n=8) | MR + MHS (n=9) |
| PW, mm | 0.82±0.025 | 0.92±0.025^c^ | 0.90±0.031^b^ |
| IVS, mm | 0.75±0.024 | 0.79±0.024 | 0.85±0.024^c^ |
| RWT | 0.41±0.008 | 0.41±0.010 | 0.40±0.012 |
| EDV, µl | 55±1.5 | 66±3.9^a^ | 80±4.8^d^ |
| ESV, µl | 21±1.4 | 28±2.1^b^ | 40±5.2^d^ |
| SV, mm | 34±0.5 | 38±2.2 | 40±2.2 |
| HR, bpm | 519±19.0 | 554±17.3 | 544±12.9 |
| CO, ml/min | 17.9±0.77 | 20.3±1.50 | 21.5±0.88 |
| **Females** |  |  |  |
| Parameters | Ctrl (n=8) | MR (n=8) | MR + MHS (n=10) |
| PW, mm | 0.79±0.012 | 0.79±0.024 | 0.80±0.013 |
| IVS, mm | 0.73±0.016 | 0.71±0.01. | 0.75±0.015 |
| RWT | 0.41±0.015 | 0.39±0.014 | 0.41±0.008 |
| EDV, µl | 51±2.3 | 51±3.1 | 50±2.1 |
| ESV, µl | 18±1.2 | 20±1.5 | 20±1.1 |
| SV, mm | 33±1.2 | 31±1.6 | 30±1.5 |
| HR, bpm | 487±22.3 | 490±18.4 | 520±14.4 |
| CO, ml/min | 16.0±0.84 | 14.9±0.78 | 15.4±0.36 |

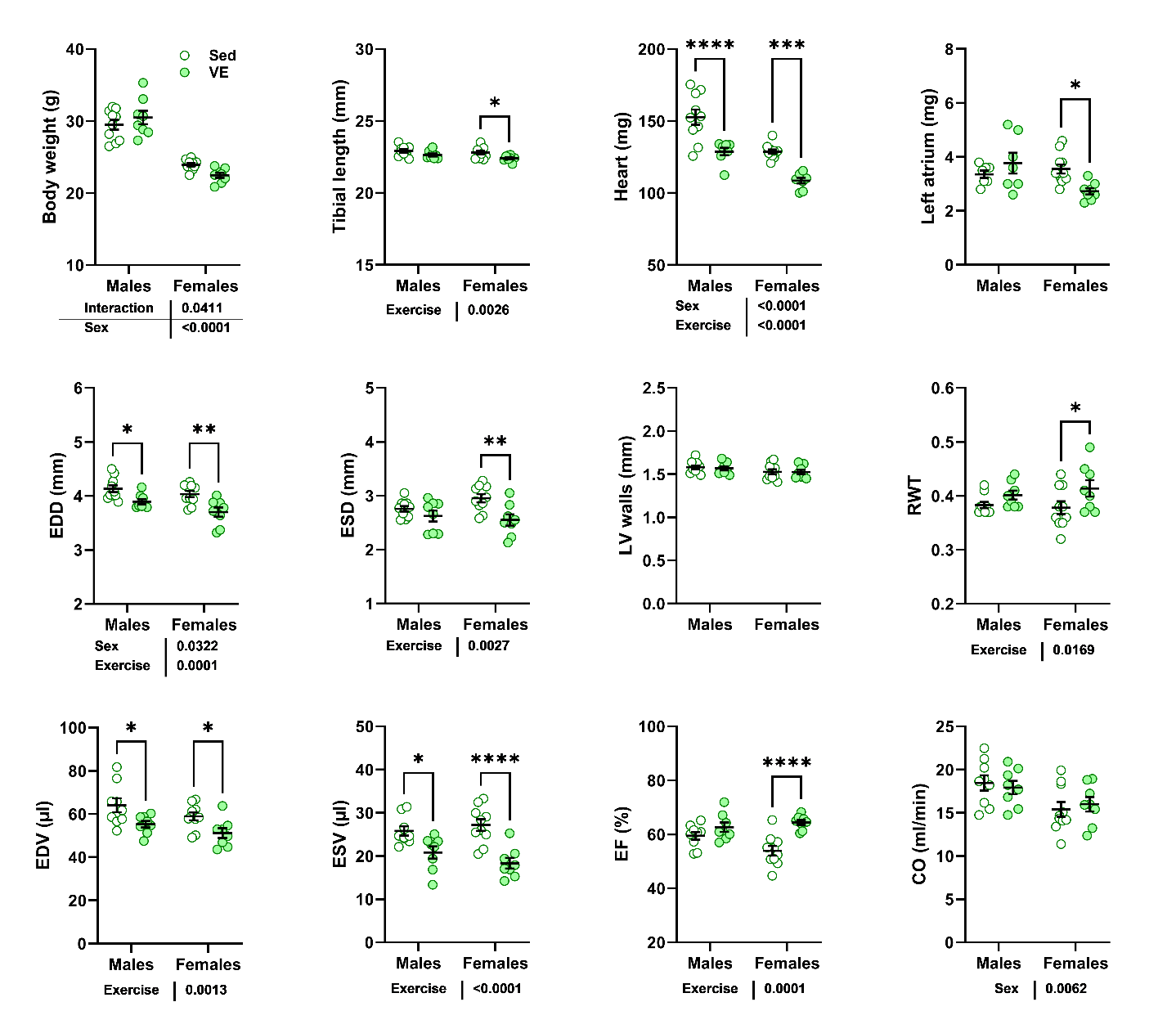

**Figure S4.** Effects of voluntary exercise (VE) for 12 weeks in 12-week-old male and female mice compared to sedentary controls (Sed). EDD: end-diastolic LV diameter, ESD: end-systolic LV diameter, LV walls: diastolic posterior wall thickness + diastolic septal wall thickness, RWT: relative wall thickness, EDV: end-diastolic volume, ESV: end-systolic volume, EF: ejection fraction, CO: cardiac output. Results are expressed as the mean ± standard error of the mean (SEM). Two-way ANOVA analysis and Holm-Sidak post-test. *: p<0.05, **: p<0.01, ***: p<0.001 and ****: p<0.0001 between indicated groups. P values below 0.05 from two-way ANOVA analysis are indicated under each graph.

**Table S5.** Influence of 12 weeks of voluntary exercise (VE) compared to sedentarity on echo parameters in male and female mice. Echo exams were performed as described in the Methods section PWd: diastolic posterior wall thickness, IVSd: diastolic inter-ventricular septum, PWs: systolic posterior wall thickness, IVSs: systolic inter-ventricular septum, EDD: end-diastolic LV diameter, ESV: end-systolic LV diameter, RWT: relative wall thickness, EDV: end-diastolic volume, ESV: end-systolic volume, SV stroke volume, HR: heart rate, EF: ejection fraction, CO: cardiac output, GLS: global longitudinal strain and GCS: global circumferential strain. Results are expressed as the mean ± standard error of the mean (SEM). P values were calculated using the Student’s T-test.

| **Males** |  |  |  |
| --- | --- | --- | --- |
| Parameters | **Sed (n=10)** | **VE (n=8)** | **P value** |
| PWd, mm | 0.83±0.012 | 0.82±0.014 | 0.79 |
| IVSd, mm | 0.76±0.014 | 0.75±0.014 | 0.28 |
| PWs, mm | 1.24±0.026 | 1.20±0.031 | 0.61 |
| IVSs, mm | 1.06±0.031 | 1.02±0.262 | 0.35 |
| EDD, mm | 4.13±0.065 | 3.89±0.048 | 0.012 |
| ESD, mm | 2.76±0.050 | 2.62±0.010 | 0.023 |
| RWT | 0.38±0.006 | 0.40±0.008 | 0.0061 |
| LV mass, mg | 122.6±4.78 | 109.0±2.99 | 0.0042 |
| EDV, µl | 66.0±3.48 | 55.2±1.50 | 0.019 |
| ESV, µl | 26.4±1.17 | 20.8±1.40 | 0.0067 |
| SV, mm | 39.6±2.63 | 34.5±0.51 | 0.11 |
| HR, bpm | 488±17.9 | 519±19.0 | 0.24 |
| EF, % | 59.6±1.28 | 62.7±0.79 | 0.17 |
| CO, ml/min | 19.1±1.00 | 17.9±0.77 | 0.39 |
| GLS | -18.9±1.37 | ---- | ---- |
| GCS | -27.6±1.86 | ---- | ---- |
| **Females** |  |  |  |
| Parameters | **Sed (n=10)** | **VE (n=8)** | **P value** |
| PWd, mm | 0.78±0.020 | 0.79±0.012 | 0.82 |
| IVSd, mm | 0.74±0.013 | 0.73±0.015 | 0.66 |
| PWs, mm | 1.08±0.031 | 1.11±0.019 | 0.43 |
| IVSs, mm | 0.99±0.031 | 0.99±0.028 | 1.00 |
| EDD, mm | 4.03±0.058 | 3.70±0.087 | 0.0044 |
| ESD, mm | 2.95±0.075 | 2.55±0.100 | 0.0053 |
| RWT | 0.38±0.012 | 0.41±0.015 | 0.072 |
| LV mass, mg | 111.0±3.02 | 96.8±3.16 | 0.0051 |
| EDV, µl | 58.9±1.84 | 51.2±2.29 | 0.0017 |
| ESV, µl | 27.2±1.35 | 18.3±0.72 | 0.00018 |
| SV, mm | 31.8±1.49 | 32.9±1.17 | 0.59 |
| HR, bpm | 484±12.8 | 487±22.5 | 0.90 |
| EF, % | 53.9±1.80 | 64.4±0.91 | 0.00022 |
| CO, ml/min | 15.4±0.84 | 16.0±0.84 | 0.63 |
| GLS | -16.1±0.81 | -16.6±1.12 | 0.71 |
| GCS | -22.5±1.37 | -27.2±0.88 | 0.020 |

**Table S6.** Influence of 12 weeks of voluntary exercise (VE) compared to sedentary on diastolic echo parameters in male and female mice. Echo exams were performed as described in the Methods section. LA diam: left atrial diameter, E: E wave. A: A wave, E slope: E wave slope, IVRT: isovolumetric relaxation time, IVCT: isovolumetric contraction time, MPI: myocardial performance index, E’: E’ wave, A’: A’ wave. Results are expressed as the mean ± standard error of the mean (SEM). P values were calculated using the Student’s T-test.

| **Males** |  |  |  |
| --- | --- | --- | --- |
| Parameters | **Sed (n=10)** | **VE (n=8)** | **P value** |
| LA diam., mm | 2.37±0.033 | 2.41±0.042 | 0.50 |
| E, mm/s | 673±29.0 | 622±26.4 | 0.23 |
| A, mm/s | 449±23.9 | 374±18.8 | 0.023 |
| E slope | -38573±2358.7 | -43919±4831.0 | 0.30 |
| E/A | 1.51±0.060 | 1.68±0.085 | 0.11 |
| IVRT, ms | 15.3±0.44 | 14.31±0.48 | 0.13 |
| IVCT, ms | 11.6±0.31 | 10.3±0.44 | 0.030 |
| MPI | 0.67±0.021 | 0.65±0.025 | 0.41 |
| E’, mm/s | -30.2±1.05 | -30.2±1.28 | 0.97 |
| A’, mm/s | -20.6±0.72 | -18.4±1.11 | 0.097 |
| E’/A’ | 1.47±0.044 | 1.66±0.052 | 0.012 |
| E/E’ | -22.6±1.28 | -22.5±0.58 | 0.29 |
| **Females** |  |  |  |
| Parameters | **Sed (n=10)** | **VE (n=8)** | **P value** |
| LA diam., mm | 2.42±0.029 | 2.17±0.024 | <0.0001 |
| E, mm/s | 601±27.49 | 566±17.6 | 0.33 |
| A, mm/s | 366±27.66 | 357±12.0 | 0.80 |
| E slope | -36807±2023.0 | -35033±1667.4 | 0.35 |
| E/A | 1.68±0.077 | 1.59±0.047 | 0.070 |
| IVRT, ms | 16.2±0.35 | 15.9±0.34 | 0.55 |
| IVCT, ms | 11.8±0.35 | 11.6±0.36 | 0.77 |
| MPI | 0.66±0.004 | 0.67±0.011 | 0.97 |
| E’, mm/s | -28.7±0.64 | -31.2±0.78 | 0.025 |
| A’, mm/s | -20.0±0.78 | -17.6±0.64 | 0.039 |
| E’/A’ | 1.45±0.005 | 1.79±0.075 | 0.00014 |
| E/E’ | -21.1±1.18 | -18.2±0.58 | 0.060 |

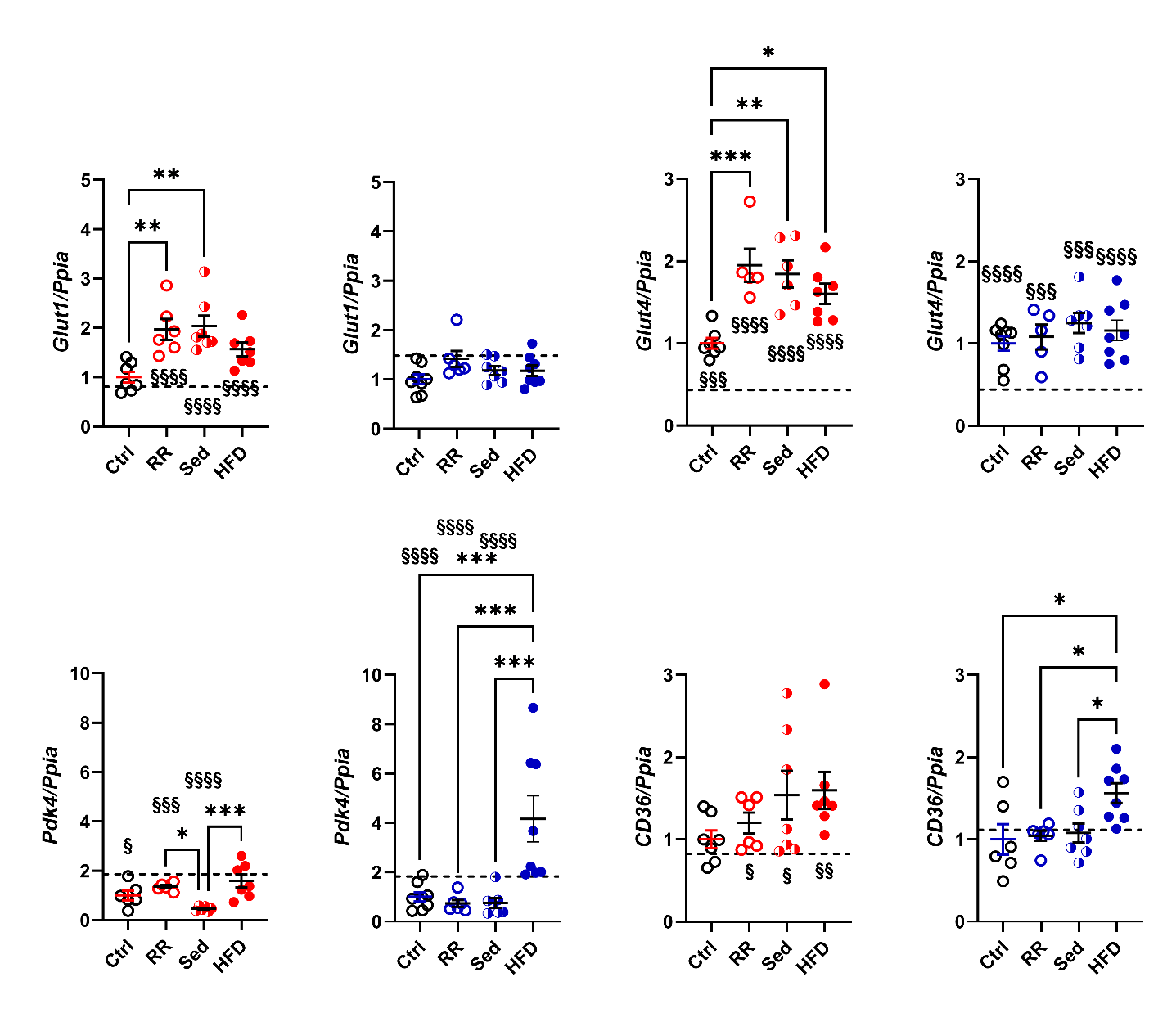

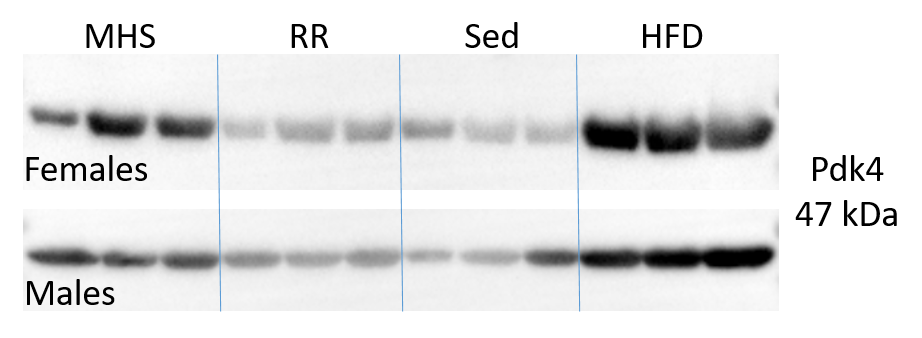

**Figure S5.** Expression levels of gene implicated in myocardial energy production after four weeks of recovery. Ctrl: control, RR: reverse remodelling group, Sed: LFD without voluntary exercise (VE) and HFD: HFD and VE. The dotted line on a graph represents the value of the indicated parameter after 28 days of MHS. The graph on the left is for females (red), and the one on the right is for males (blue). One-way ANOVA followed by Holm-Sidak post-test. *: p<0.05, **: p<0.01, ***: p<0.001 and ****: p<0.0001 between indicated groups. §: p<0.05, §§: p<0.01, §§§: p<0.001 and §§§§: p<0.0001 between the indicated group and MHS group using the Student’s T-test. Western blots of Pdk4 after MHS and 4 weeks of recovery (RR, Sed and HFD).
